# ABC-seq expands small RNAs content with randomized adapter pools operation

**DOI:** 10.1101/2024.08.30.610480

**Authors:** Huahang Yu, Mengying Ye, Yingying Guo, Yating Sun, Ke Cao, Dexin zhang, Feng Chen, Yongxi Zhao

## Abstract

Cellular gene expression is widely regulated by a large number of small RNAs (smRNAs). The ability to in-depth analysis of smRNAs content is vital for comprehensive understanding their architecture and function. However, experimental approaches capable of realizing both high-content and low-byproduct sequencing analysis are lacking. Here, we develop ABC-seq (randomized <u>A</u>dapter pool dimer <u>B</u>locking and <u>C</u>leavage in smRNA sequencing). This method designs randomized adapter pools to recognize more smRNA species. Moreover, it operates two product structure-differentiated enzymatic reactions to maximally eliminate the byproducts of adapter dimers. Using ABC-seq, we detected a broader range of smRNAs including miRNAs, scRNAs, and snoRNAs in rat hypertrophic cardiomyocytes. Upon further analysis of miRNAs, we detected 66 more miRNAs compared to the control method, providing new insights into smRNA-directed gene regulation mechanisms. Furthermore, we revealed cancer drug response -associated smRNAs and uncovered their pivotal role in immune response modulation in non-small cell lung cancer. ABC-seq represents an evolutionary strategy for reshaping the smRNA landscape boundaries and paves the way for a deeper understanding of complex gene regulation networks.

## Introduction

Cellular processes and signal pathways have been confirmed to be widely regulated by a vast range of non-coding RNAs (ncRNAs) in both physiological and pathological contexts^1^. Among different ncRNA classes, small RNAs (smRNAs) are known as regulatory ncRNAs with short sequences (<200 nt). The main smRNAs include microRNAs (miRNAs), PIWI-interacting RNAs (piRNAs), small nuclear (snRNAs) and nucleolar (snoRNAs). They perform complex regulatory roles in many human diseases^2–5^. Especially, their aberrant expression are involved in cardiac hypertrophy and cancer drug response^3,5–7^. Comprehensive and extensive analysis of the expression of genome-wide smRNAs holds broad significance for understanding their architecture, regulation function and mechanism^4^.

High-throughput next-generation sequencing of small RNA (smRNA-seq) is the standard method to detect the diversity of small RNAs at the transcriptome-wide level^6^. Before sequencing reaction, target RNA molecules are transformed into sequence libraries. These sequences contain different target sequences in the middle, with the same 5’ adapter and 3’ adapter at the ends. Adaptors ligation is the dominant strategy of library construction for smRNA-seq^2,4,6,8–25^. In this strategy, two single-stranded (ss) adaptors is successively ligated to the 3’ hydroxyl end and 5’ phosphate end of smRNAs by two different ssRNA ligases^2,6,8–10,13,16–18,20^. The various sequences and structures of smRNA cause significant difference in adapter ligation efficiency^6,8,22,26,27^. Strategies involves the ligation of adaptors in a predetermined orientation to the ends of RNAs are generally in capturing a wide range of small RNA species, including those with diverse sequences and structures^6,8,22,26,27^. The use of improved ligation adapters, such as the randomized adapters (NNN adapters) with several random nucleotides at the ligation ends, reduced the bias on the order of 100-fold and increased the sequence coverage compared to non-randomized adapters (non NNN adapters) ^13,14,22,24–26,28,29^. However, a significant challenge arises during sequencing library preparation due to the formation of large quantities of adapter dimers. These byproducts can compete with smRNA for adapter ligation efficiency and have a higher likelihood of preferential amplification in subsequent PCR, leading to low detected smRNA content. There were several strategies that can reducing adapter dimers^2,6,8–10,16,20,27,30,31^. Nevertheless, applying these approaches to smRNA-seq with randomized adapter has been difficult because the reduction of adapter dimer relies on hybridizing excess adapters or adapter dimers with known sequences, which cannot be applied to randomized adapters. Thus, a high-content and low-byproduct sequencing method for smRNAs remains a critical need.

Here we developed a smRNA-seq method called ABC-seq (randomized <u>A</u>dapter pool dimer <u>B</u>locking and <u>C</u>leavage in smRNA-seq). By employing two low-bias randomized adapter pools, ABC-seq maximizes detected smRNA content, regardless of secondary structures of these RNAs. Moreover, we operate two product structure-differentiated enzymatic reactions. One is inhibiting the formation of adapter dimers, and another one is cleaving adapter dimers after their formation. This operation allowed us to minimize the byproducts of adapter dimers. Appling ABC-seq to rat hypertrophic cardiomyocytes, we detected a greater variety of smRNAs, including miRNAs, scRNAs, snoRNAs and fewer rRNAs. Especially, we detected 66 more miRNAs compared to the negative control method. Additionally, we detected 107 miRNAs over an 853-fold range associated non-small cell lung cancer (NSCLC) and revealed that smRNAs play pivotal role in immune response modulation. Overall, ABC-seq addresses the challenge of reducing randomized adapter dimer in smRNA-seq. This advancement may represent a significant upgrade in smRNA-seq and pave the way for more accurate and comprehensive smRNA profiling.

## Results

We developed ABC-seq to simultaneously reduce adapter dimer byproducts and increase smRNAs content. In this method, we introduced three molecular designs (Fig.1). First, two randomized adapter pools are employed to overcome the sequence bias and recognize more RNA classes during two ligation reactions. Moreover, two product structure-differentiated enzymatic reactions are designed to operate randomized adapter pools: block the formation of adapter dimer and cleave the byproduct of adapter dimer. In brief, those randomized adapter pools are designed with 5’ NNN end and 3’ NNNNN end (N=A, T,C or G), respectively. They contain more than 64-type terminal sequences to widely enhance ligation efficiency for diverse smRNAs. On the other hand, adapter dimers can be generated by direct ligation of two randomized adapter pools. These byproducts compete with smRNAs in both ligation process and sequencing process, leading to significantly low detected content of smRNAs.

**Fig 1:**
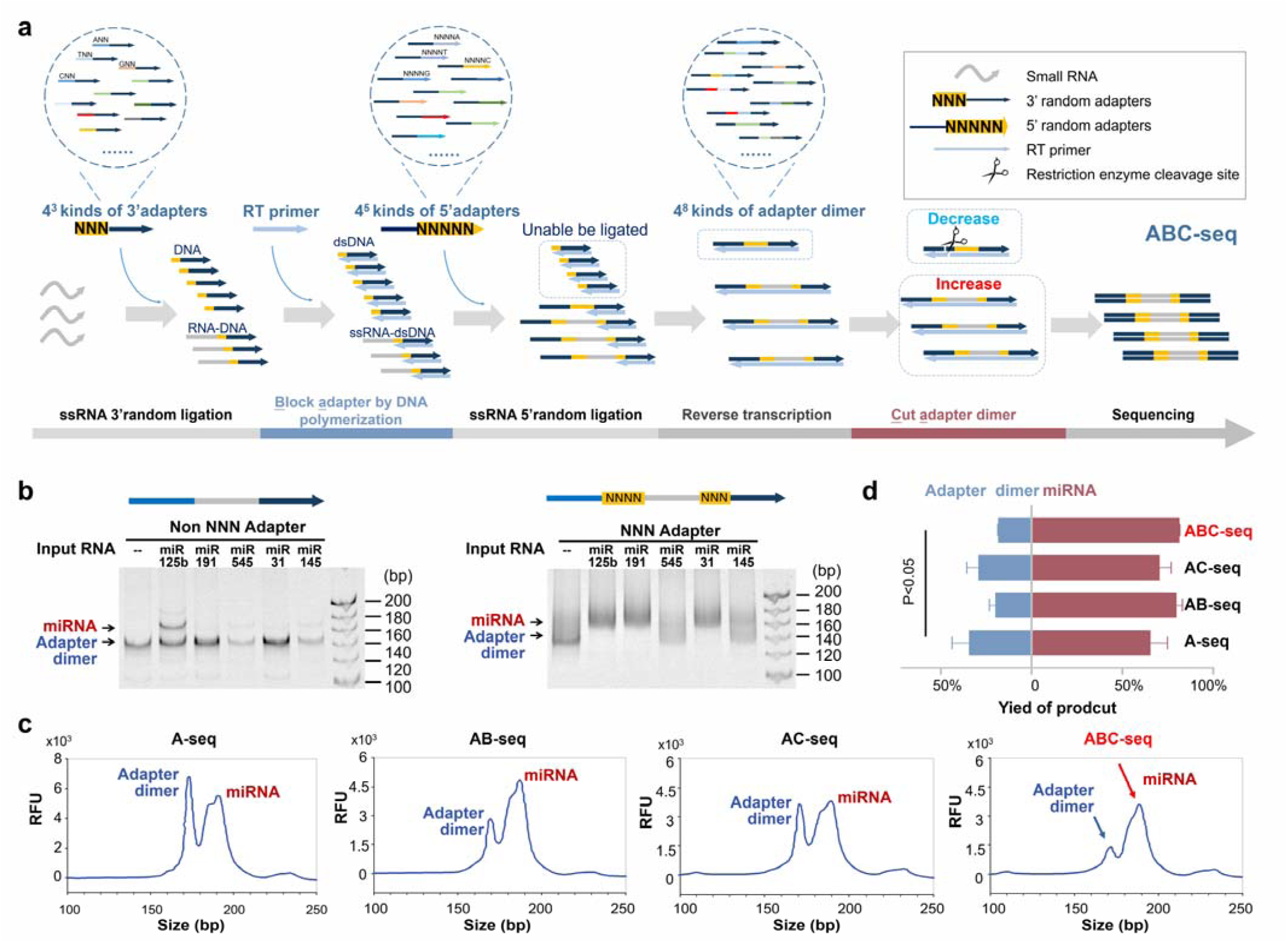
Overview of ABC-seq. **a** Schematic presentation of the ABC-seq technology. **b** Gel electrophoresis demonstrating the reduction of ligation bias by using randomized adapters (NNN adapters). Left panel: libraries constructed without randomized adapters (non NNN adapter) for five different secondary structures of miRNA and the negative control. Right panel: libraries constructed with pooled adapters (NNN adapter) for the same miRNAs and the negative control. **c** Bioanalyzer traces of libraries prepared with 100 ng small RNA using A-seq, AB-seq, AC-seq, and ABC-seq. The left peak represents the position of the adapter dimer, and the right peak represents the position of the miRNA ligation product. **d** Statistical analysis of the peak area from panel (c). Reductions of adapter dimer in A-seq and ABC-seq were significant (K-S test p-value <0.05). Quantification of the miRNA ligation products and adapter dimer, with their combined total representing 100% of the sample (n=3 in each group).

For blocking the formation of adapter dimer, we design a RNA template-stopped DNA polymerization strategy. After the first ligation (3’ randomized adapter to the 3’ hydroxyl end of smRNAs), excess 3’ randomized adapters can compete with the ligation product of smRNA-3’ randomized adapter in the second ligation (5’ randomized adapter to the 5’ phosphate end of smRNAs). One reasonable solution is to transform the 3’ randomized adapters (ssDNA) into dsDNA, while retaining the single-stranded structure of smRNAs. A DNA polymerase without the ability of using RNA template (e.g., Phi 29 DNA polymerase) is utilized to catalyze this polymerization reaction. During this polymerization reaction, the excess 3’ randomized adapters can be formed dsDNA by DNA extension to their 5’ NNN ends. In contrast, the DNA extension is stopped at RNA templates in the ligation product of smRNA-3’ randomized adapter. Then the dsDNA can not be ligated by the ssRNA ligase during in the second ligation. In this way, the formation of adapter dimer may be maximally blocked, mainly dependent on both DNA polymerase efficiency and ssRNA ligase fidelity.

For cleaving the byproduct of adapter dimer, we design 5’-DNA-RNA-3’ chimeras rather than tranditional ssRNAs as the 5’ randomized adapter pools. This chimera design result in the adapter dimer as 5’-DNA-RNA-NNNNNNNN-DNA-3’ and the smRNA ligation product as 5’-DNA-RNA-smRNA- NNNNNNNN-DNA-3’. As we know, the -N…N- sequences can not be recognized by certain DNA restriction enzyme (RE). Therefore, we require specific types of DNA RE that recognized 3’ downstream DNA sequences but cleave 5’ upstream DNA position with an appropriate sequence length. Using this enzymatic cleavage reaction, the adapter dimer can be degraded after reverse transcription reaction, whereas the smRNA ligation product is preserved for subsequent amplification and sequencing. In this way, the potential adapter dimer may be eliminated. Overall, our ABC-seq employs above-mentioned two enzymatic designs to maximally reduce adapter dimer byproducts while increasing smRNAs content.

When only using the blocking design or the cleaving design, the method is referred to as AB-seq or AC-seq, respectively. The negative control using neither design is referred to as A-seq.

We then applied ABC-seq to sequence small RNAs in H9C2 cells and compared the results with those obtained using A-seq (Fig. S2). Notably, the bioanalyzer results showed that AB-seq, AC-seq, and ABC- seq significantly reduced adapter dimers compared to A-seq (Fig. 1c, d). Specifically, the proportion of adapter dimers in small RNA sequencing decreased from 20.29% to 5.29% in ABC-seq, demonstrating the effectiveness of the adapter pool operation in minimizing byproduct and improving sequencing quality.

### AB-seq limits the formation of randomized adapter dimer

To evaluate the necessity of adapter pool, we constructed libraries using both NNN and non-NNN adapters for five representative miRNAs with distinct secondary structures (Fig. 1b, S1). We designed 3 random nucleotides at the 5’ end of 3’ adapters and 5 random nucleotides at 3’ end of 5’ adapters. The results demonstrated that NNN adapters significantly reduced ligation bias, resisted with previous researches. However, we found that adapter dimers in the control were also increased. We reasoned that it is difficult to blunt the terminal random nucleotides of the 3’ adapters due to hybridization issues with the RT primer (Fig. 1b).

We observed the nucleotide differences between the products of miRNA 3’ adapter ligation and excess 3’ adapters. Specifically, the product of miRNA 3’ adapter ligation exhibited RNA nucleotides at its 5’ end, whereas the excess 3’ adapter contained DNA nucleotides at its 5’ end. Additionally, the 5’ end of the excess 3’ adapter consists of random nucleotides. The RT primer bearing the specified sequence failed to hybridize with the NNN end, resulting in the formation of adapter dimers.

Therefore, we tested five methods to block excessed 3’ NNN adapters (Fig. 2b, 2c). For method II, the overhang terminal random nucleotides of NNN RT primer prevent the reverse transcription. For method III, the low hybrid efficiency of the NNN RT-primer and the NNN adapter results in inefficient ligation of 3’ adapter and 5’ adapter. However, the target product did not significantly increase, possibly due to inefficient reverse transcription caused by the overhang of the RT primer. For method IV, there are still adapter dimer formed in the control, suggesting that the RT primer was hard to block NNN adapter. For method V, the 3’ NNN adapter was filled to double strand by RNA template-stopped DNA polymerase while the ligation of smRNA and 3’ adapter can keep 5’ overhang be single strand. The result reveals that RNA template-stopped polymerase blocking performed better than other ways, suggesting that the DNA polymerase is helpful for the blunt of 3’ adapter.

**Fig. 2:**
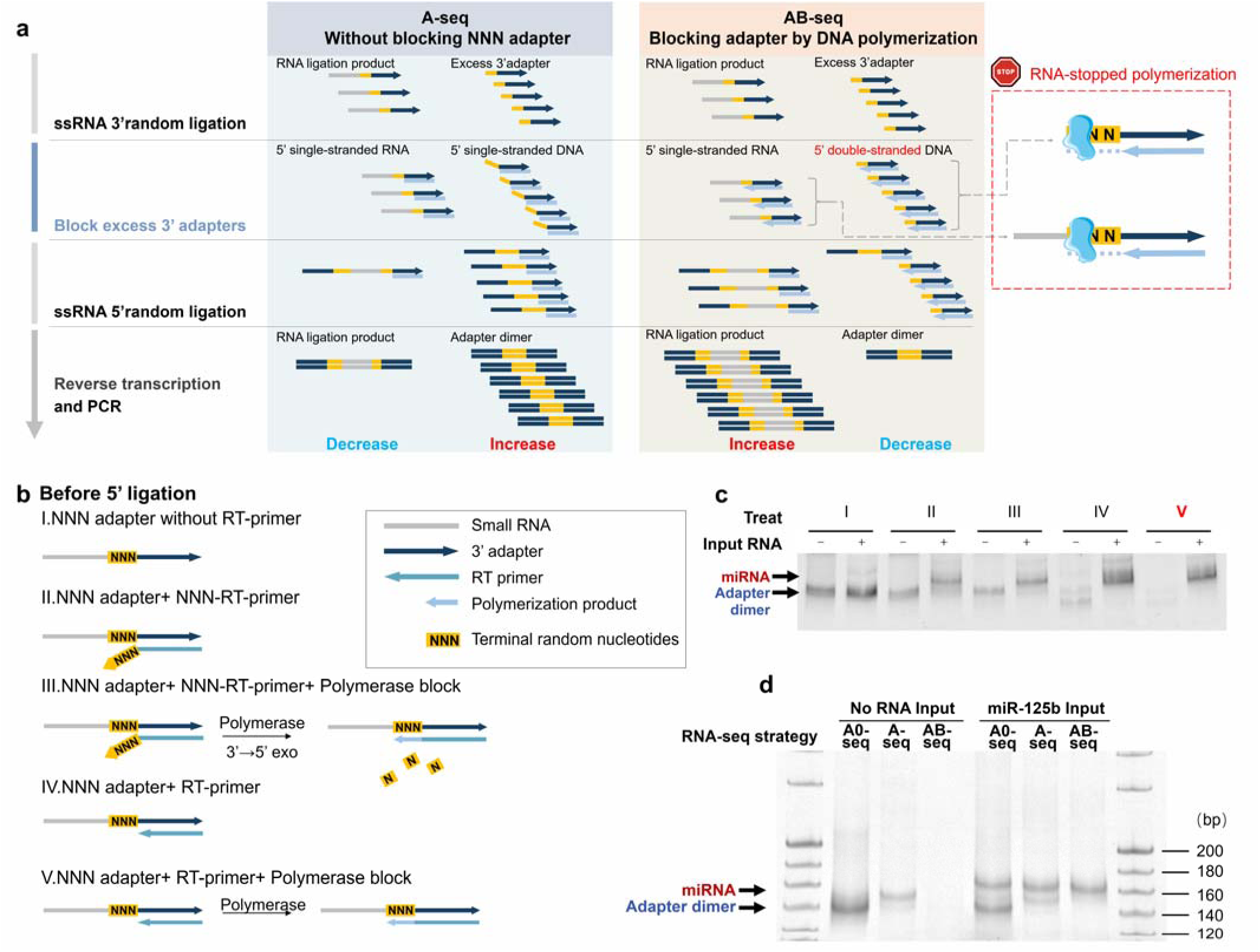
AB-seq limits the formation of adapter dimer. **a** Schematic diagram illustrating the principle of AB-seq. **b,c** Schematic diagrams and polyacrylamide gel electrophoresis(PAGE) characterizing the use of different RT primer blocking strategies to blunt excess 3’ adapters. Method V shows the lowest adapter dimer content. **d** Comparison of the sequencing library constructed by A0-seq(A-seq without any blocking strategy), A-seq and AB-seq. 10 ng miR-125b was used as input and the PCR amplification products were analyzed by 10% PAGE.

Based on this result, AB-seq was designed to inhibit the formation of NNN adapter dimers (Fig. 2a, 2d). We reasoned that it should be possible to prevent <u>a</u>dapter dimer formation by <u>b</u>locking the excess 3’ adapter sticky ends (AB-seq). Phi29 DNA polymerase, an RNA template-insensitive DNA polymerase, was selected to blunt the excess 3’ adapter before 5’ ligation (Fig. 2c). Notably, we also observed that adapter dimer was still formed in AB-seq (Fig. 1c). One explanation is that the purchased polymerase may have non-ideal efficiency in filling 5’ overhang of dsDNA, leading to the incomplete blunt end on the excess 3’ adapters. The other explanation is that the purchased RNA ligase may perform undetected feature that it can ligate ssRNA to dsDNA. These explanations both confirm the probable formation of adapter dimers. Moreover, there were multiple rounds round of PCR amplification in AB-seq when using low-input RNA samples, which amplified the few adapter dimers remained in AB-seq.

### AC-seq reduces randomized adapter dimer after generation

To eliminate adapter dimers formed after generation, we developed AC-seq. Traditional Type II restriction enzymes cleave inside the recognition sequence and cannot reduce the pool of 4^8^ types of adapter dimers all at once (Fig. 3a). We noticed that the adapter dimer lacked smRNA insert (18-30 bp) compared to the smRNA library product. Indeed, all of adapter dimers share a common structural feature, which includes 8 random nucleotides in the middle, flanked by known nucleotide regions (Fig. 3a).

**Fig 3:**
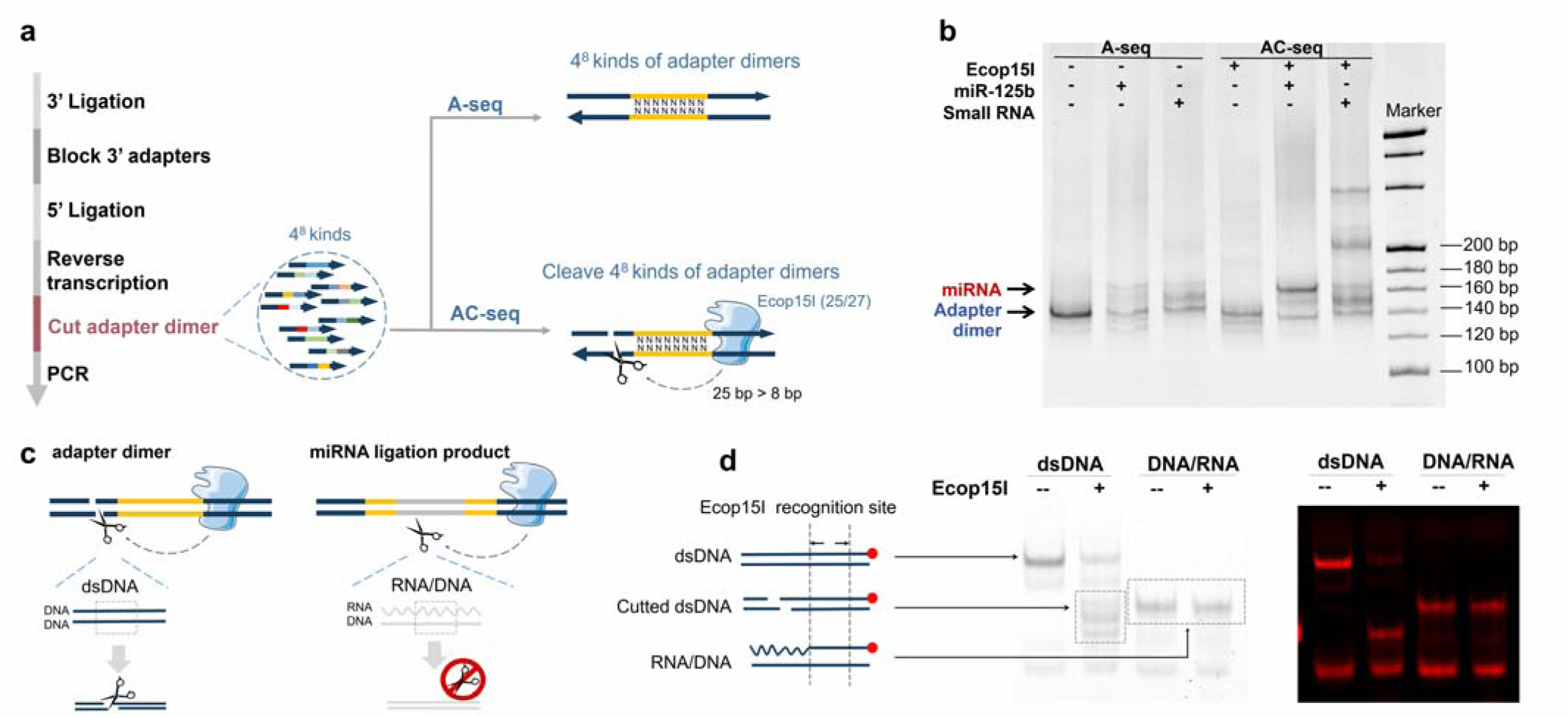
AC-seq depletes adapter dimer after formation. **a** Schematic diagram illustrating the principle of AC-seq. AC-seq uses Type III RE (EcoP15I), which cut outside the recognition sites. The enzyme bypasses the random nucleotide regions for cutting, which allows for general cleavage of the entire 4^8^ kinds of adapter dimers. **b** Comparison of library construction efficiency between AC-seq and A-seq using 10 ng miR-125b and 200 ng small RNA as input. PCR amplification products were analyzed by 10% PAGE. **c** Specificity of AC-seq in cutting adapter. Adapter dimers at the cutting site of TypeIII RE are dsDNA, fitting the enzyme’s conditions. miRNA ligation products of TypeIII RE are RNA/DNA hybrids, which do not fit the enzyme’s conditions. **d** The EcoP15I enzyme cuts dsDNA but not RNA/DNA hybrids. Left panel: Gel electrophoresis imaged under UV light at 254 nm. Right panel: Gel electrophoresis imaged under 650 nm excitation light. Cy5 fluorescence is indicated by red dots, solid blue lines represent DNA oligos, and dashed blue lines represent RNA.

However, the target product contains an unknown nucleotide region of at least 26 bp, which differs from adapter dimers. Therefore, we hypothesized that the adapter dimer could be cleaved by REs that can cut across the 8 random nucleotides region.

We first tested Type IIS RE in AC-seq, which recognize 6-12 consecutive bases and cleave within recognition sequence (Fig.S3). Firstly, MwoI was chosen due to the random sequence inside the recognition sequence as long as 7 nt. We performed AC-seq on 1ug miR-125b and 10ng miR-125b. The result shows that adapter dimer was reduced while the smRNA ligation product proportional increased. That’s because both adapter dimer and smRNA ligation product can be amplified during PCR amplification. The decrease of adapter dimer leads to the preferential amplification of smRNA ligation product. We further tested our workflow in 200 ng Hela cell small RNA (Fig.S3). However, there is a little amount of smRNA ligation product in the gel. Prior work has demonstrated that the number of RNA nucleotides proximate to the end of nucleic acid molecule can influence RNA ligase enzyme ligation. We reasoned that the recognition sequence of MwoI may limit the design of 5’ adapters, which decrease the ligation efficiency of RNA ligase enzyme.

Based on this finding, we optimized 5’ adapter design, which is adding 20 RNA nucleotides at 3’ end. However, the increase of RNA nucleotides limits the choose of TypeIIS REs due to the length of their recognition sequence maximized to only 11 nt, which is far shorter than 20 nt. So we further upgrade AC-seq by using TypeIII RE (Ecop15I), which cleaves downstream of the recognition site, typically 25-28 bases away (Fig.3a). Due to the amino acids of them are partitioned into two separate domains, linked by a short polypeptide connector.

Subsequently, we present the principles and outcomes of employing Type III REs for adapter dimer cleavage in Fig. 3c. The key to this strategy lies in the use of Type III REs, which are DNA restriction endonucleases. We found that it can only cleave the dsDNA rather than RNA /cDNA hybrid, consisted with our hypothesis (Fig. 3c, 3d). We observed that the adapter dimers precisely meet the specific conditions required for the activity of this particular restriction endonuclease, whereas the miRNA ligation products do not. Consequently, we hypothesized that this restriction endonuclease could selectively excise adapter dimers. As demonstrated in Fig. 3b, the enzyme effectively cleaves the adapter dimers, leading to their significant reduction. Concurrently, we observed an increase in the yield of the desired ligation products. This supports the hypothesis that reducing the presence of adapter dimers enhances the amplification efficiency of miRNAs in the library, consistent with previous findings.

Moreover, Type III REs are infrequently used in molecular biology. Our strategy, however, takes advantage of their special features to achieve to meet the need for long distances between recognition sites and cutting sites, expanding the use of Type III REs in the field of nucleic acid sequencing.

### ABC-seq profiles comprehensive smRNA landscape in hypertrophic cardiomyocytes

To determine the efficacy of different sequencing strategies (A-seq, AC-seq, AB-seq, and ABC-seq) in detecting high content smRNAs, we performed these four methods in hypertrophic cardiomyocytes.

ABC-seq offered a higher single-molecule RNA capture efficiency compared to A-seq (P < 0.0001) (Fig.2a). The correlation analysis indicated that A-seq and AC-seq exhibited higher correlation with each other, whereas AB-seq and ABC-seq showed a stronger correlation (ρ=0.95, Fig. 4b). This may be because the removal of adapter dimers primarily enhances the amplification efficiency of the miRNA library without necessarily increasing the yield of target products. In contrast, AB-seq, which incorporates a critical step during the ligation reaction, not only reduces the formation of adapter dimers but also facilitates the production of more target miRNA products.

**Fig 4:**
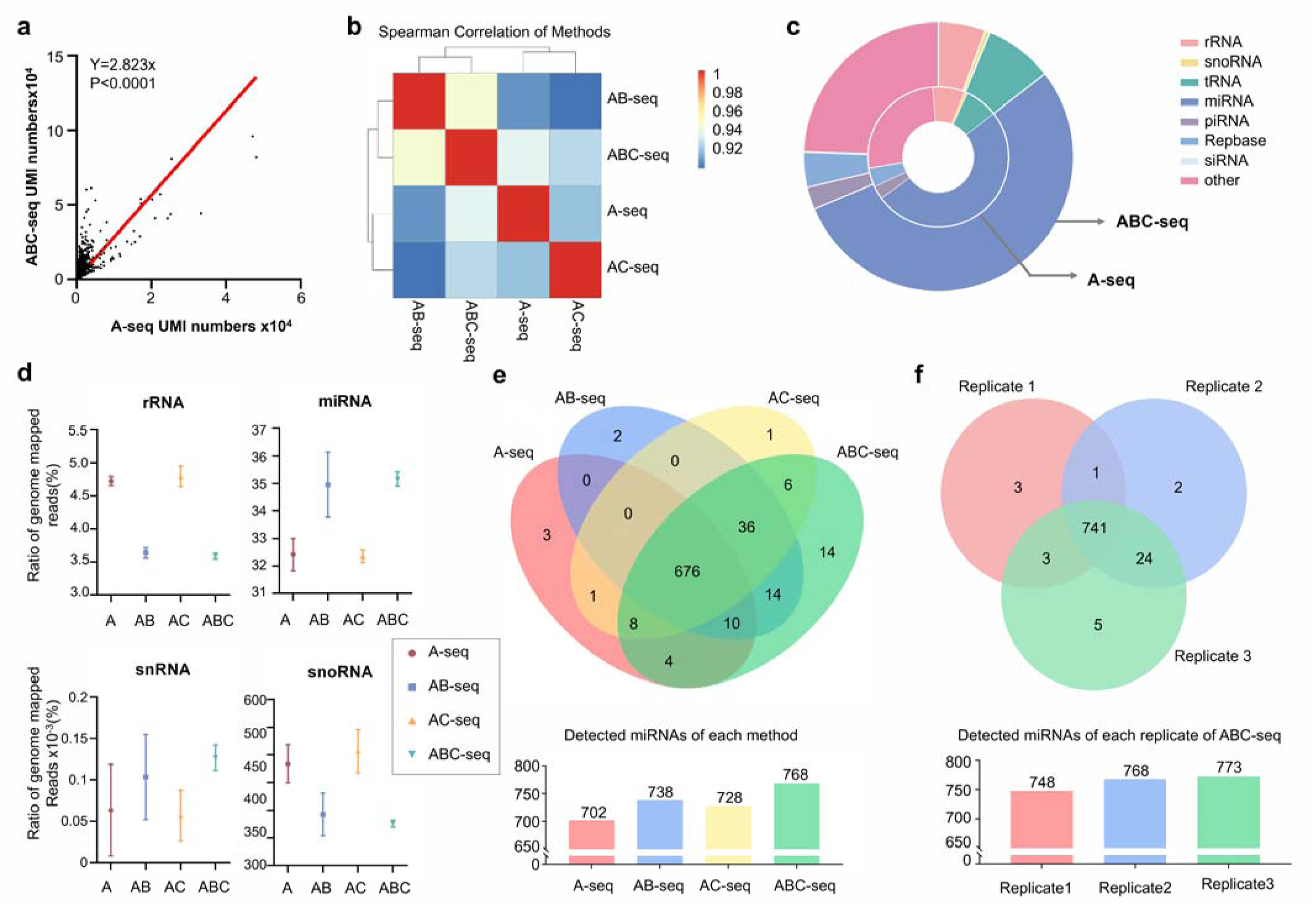
Comparative analysis of smRNA-seq methods. **a** Correlation of RNA level quantified by UMI numbers between ABC-seq and A-seq. Each dot corresponds to one gene. Pvalue means that the relationship represented by the slope is statistically significant. **b** Comparison of A-seq, AB-seq, AC-seq, and ABC-seq using Spearman correlation analysis. **c** Distribution of different types of smRNAs in hypertrophic cardiomyocytes detected by ABC-seq or A- seq (n = 3). **d** Comparison of the genome-mapped read ratios of rRNA, miRNA, snRNA, and snoRNA detected by A-seq, AB-seq, AC-seq, and ABC-seq (n = 3). **e** miRNAs detected by A-seq, AB-seq, AC- seq, and ABC-seq (100 ng small RNA input) are plotted as Venn Diagrams and bar chart, with RNAs from hypertrophic cardiomyocytes. Numbers of detected miRNA by 4 methods are listed. The diagram at the bottom of Figure 4d shows the number of miRNAs detected across different combinations of methods, including those common to all four methods, three methods, two methods, and those detected by only one method. **f** miRNAs detected in three replicates of ABC-seq (100 ng small RNA input) are plotted as Venn diagrams and bar chart, with RNAs from hypertrophic cardiomyocytes . Numbers of detected miRNA by three replicates are listed. The diagram at the bottom of Figure 4e shows the number of miRNAs detected across replicates, including those common to three replicates, two replicates, and those detected by only one replicate. This analysis demonstrates the reproducibility of the ABC-seq method, with a high number of miRNAs consistently detected across all three replicates, indicating reliable and consistent performance.

Next, we performed four smRNA-seq methods on smRNAs extracted from hypertrophic cardiomyocytes and compared the different classes of detected smRNAs. ABC-seq demonstrated a superior capability for detecting a broader range of miRNAs and snRNAs compared to the other methods (Fig. 4c, 4d). This enhanced detection might be attributed to the randomized adapter pooling strategy employed in ABC-seq, which could reduce biases associated with adapter ligation, thereby allowing for a more comprehensive capture of these RNA species. To further compare the methods, we used Venn diagrams to illustrate the overlap and uniqueness of miRNAs identified by each method. The Venn diagram in Figure 4e demonstrates that ABC-seq identifies a larger number of unique miRNAs compared to AC-seq, AB-seq, and A-seq, highlighting its superior sensitivity. Specifically, ABC-seq detected 66 more miRNAs compared to A-seq. Additionally, we used Venn diagrams to assess the reproducibility of each method (Fig.4f). ABC-seq showed a higher degree of overlap in detected miRNAs across replicates, indicating greater reproducibility compared to the other methods. Additionally, ABC-seq detected a significantly lower proportion of rRNA, suggesting higher specificity of the method for small RNA species of interest.

Interestingly, AC-seq was found to detect a higher number of snoRNAs, whereas both AB-seq and ABC-seq showed lower detection rates for this class of RNAs. One possible explanation for this discrepancy could be the influence of secondary structures in snoRNAs. The addition of phi29 polymerase in AB-seq and ABC-seq might alter the snoRNA structure due to its 3’ -5’ exonuclease or polymerase activities. This structural alteration could negatively impact the ligation efficiency of the adapters, leading to reduced snoRNA detection, which is consistent with prior research. Another explanation is that AC-seq, which relies on the byproduct of enzymatic cleavage, may favor the amplification of low-abundant RNAs like snoRNAs, thereby improving their detection. This finding supports the notion that snoRNAs possess complex secondary structures distinct from other ncRNAs, consistent with previous studies.

These findings suggest that ABC-seq is more advantageous for detecting miRNAs and snRNAs, which do not have the same complex secondary structures as snoRNAs. In contrast, AC-seq appears to be more effective for capturing snoRNAs and other RNAs with unique structural features. The differences in detection underscore the importance of selecting the appropriate sequencing method based on the specific small RNA species of interest.

### ABC-seq reveals the smRNA networks in cardiac hypertrophy

Non-coding RNAs (ncRNAs), particularly cardiac-specific small RNAs, play unique regulatory roles in both physiological and pathological cardiac hypertrophy (CH)^32,33^. Angiotensin II (Ang-II) is a critical factor in mediating CH. Multiple signaling pathways have been shown to be involved in mediating Ang- II–induced CH, including the G-protein–coupled Ang-II type-1 receptor (AT1R) signaling, Ang-II/AT1R activates Ca2+/calmodulin/calcineurin signaling, and Ang-II/AT1R activates transforming growth factor- β expression and increases SRF-mediated gene expression. To gain a more comprehensive understanding of the signaling pathways involved in the induction process, we employed the ABC-seq on total RNA from hypertrophic and normal cardiomyocytes. We mediated cardiomyocytes to hypertrophic cardiomyocytes by AngII. (Fig. 5a-c). The sequencing reads revealed distinct miRNA length distribution patterns between the two cell types. In normal cardiomyocytes, the miRNA length distribution was more uniform, whereas hypertrophic cardiomyocytes exhibited a higher proportion of 20-nucleotide fragments (Fig. 5d). This shift in miRNA length distribution suggests potential alterations in miRNA processing or stability in hypertrophic cells.

**Fig 5:**
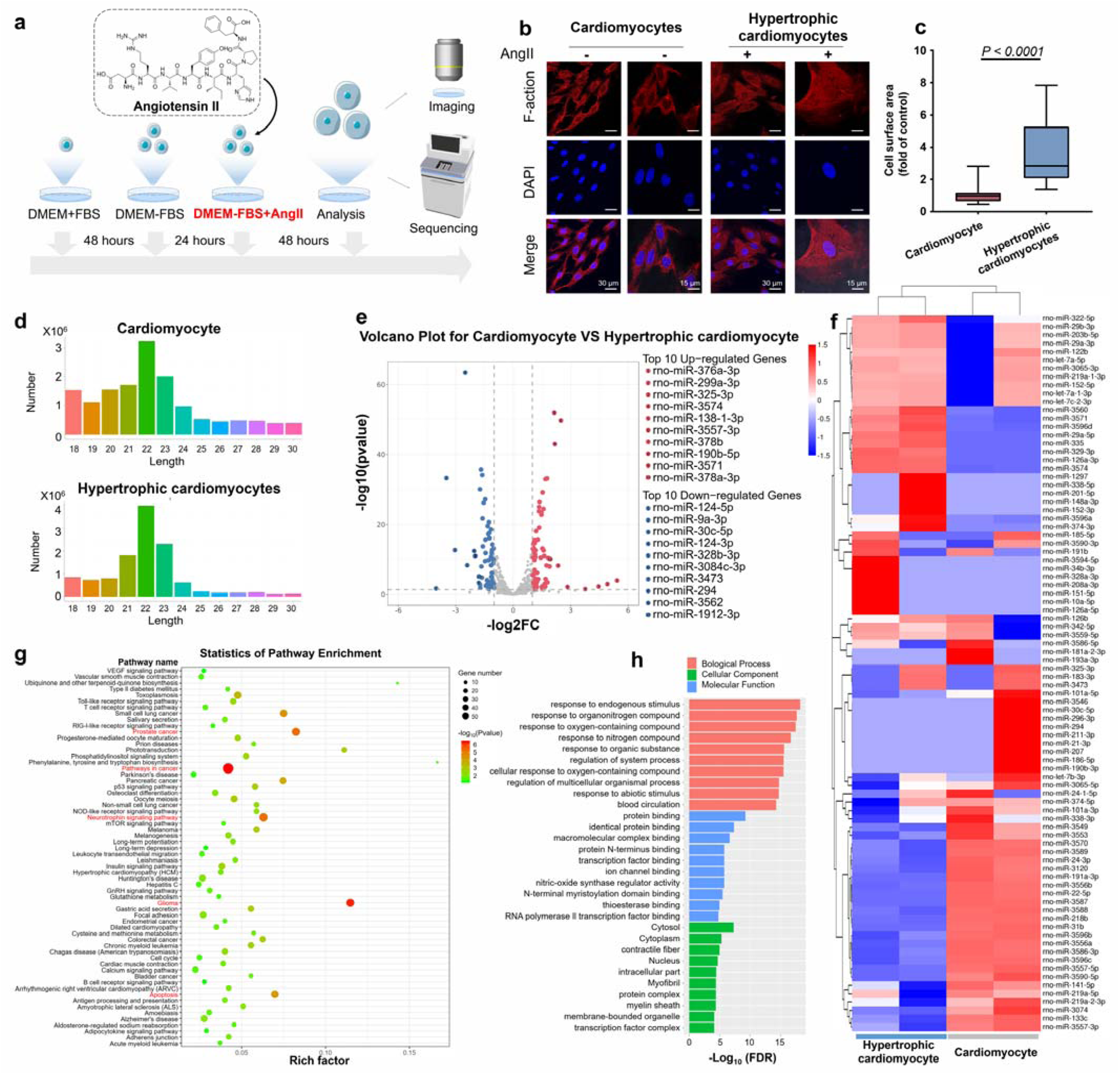
Analysis of miRNAs in Cardiomyocytes and Hypertrophic Cardiomyocytes Using ABC-seq. **a** Schematic diagram of the culture process for inducing cardiomyocyte hypertrophy. **b** Imaging of cardiomyocytes and hypertrophic cardiomyocytes. Hypertrophy was assessed on the basis of the level of sarcomere organization. Scale bar, 30 μm. **c** Quantitative analysis of cell surface area (n = 20 independent experiments, Paired t-test, p < 0.05). **d** Length distribution of miRNAs in cardiomyocytes and hypertrophic. **e** Volcano plot of differentially expressed miRNAs, showing upregulated and downregulated miRNAs, with selection criteria of |log2(FC)| ≥ 1.00 and FDR ≤ 0.01. **f** Heatmap of differential miRNA expression analysis between cardiomyocytes and hypertrophic cardiomyocytes, with selection criteria of |log2(FC)| ≥ 1.00 and FDR ≤ 0.01 (n = 2). **g** KEGG pathway enrichment analysis of significantly downregulated miRNA target genes. Each row repress ents an enriched pathway, with bubble color reflecting significance (green to red, increasingly significant), bubble size indicating overlap between differentially expressed genes and pathway genes, and the horizontal axis representing the Rich Factor, which considers both the overlap and the total number of genes in the pathway. Higher values indicate greater enrichment of differentially expressed genes in the pathway. **h** GO enrichment analysis of target genes of differentially downregulated miRNAs between hypertrophic cardiomyocytes and cardiomyocytes. The 10 most significant GO terms in each GO category are shown, with different colors representing different GO categories. The length of each bar indicates the significance of enrichment, with longer bars representing greater significance.

To further understand the molecular changes, we conducted differential expression analysis between normal cardiomyocytes and hypertrophic cardiomyocytes. We detected 131 miRNAs differentially expressed (|log2(FC)| ≥ 1.00, p < 0.05) between normal and hypertrophic cardiomyocytes. The analysis revealed that 61 miRNAs were upregulated and 70 miRNAs were downregulated in hypertrophic cardiomyocytes compared to normal cardiomyocytes (Fig. 5e). Furthmore, we performed hierarchical clustering analysis to compare these differentially expressed miRNAs, whichshowed significant expression patterns between the two conditions (Fig. 5f). Consistent with previous studies, we observed that miR-27a-5p and miR-146a were markedly upregulated in the hypertrophic cardiomyocytes especially in those CH model mediated by Ang-II^34,35^. Conversely, miR-26b-3p and miR-30c were significantly downregulated in the hypertrophic model, aligning with earlier findings^36,37^.Further functional enrichment analysis revealed that the target genes of these differentially expressed miRNAs are implicated in several key signaling pathways, including pathways in cancer, prostate cancer, neurotrophin signaling pathway, glioma, and apoptosis (Fig. 5g, 5h, S4-7). This analysis suggests a complex interplay between these miRNAs and the molecular mechanisms driving hypertrophic growth.

### ABC-seq profiles the smRNA landscape in immune cells of NSCLC and its association with response to chemoimmunotherapy

Recently, RNA-seq analysis of cancerous cells and immune cells from non-small cell lung cancer (NSCLC) patients revealed a correlation between cancer therapies and the immunosuppressive T cell state^38^. Given the crucial role of small RNAs in regulating immune responses, we utilized ABC-seq to analyze small RNA in CD4+ T cells from the peripheral blood of NSCLC patients. It aimed to uncover the effects of chemotherapy and immunotherapy on NSCLC. Peripheral blood samples were collected from three groups: treated non-small cell lung cancer (NSCLC) patients (n=6), treatment-naive NSCLC patients (n=6), and healthy individuals (n=6) (Figure 6a). In total, we identified 790 miRNAs, with 107 of them being differentially expressed across groups (|log2(FC)|≥1.00; P value≤0.05). Then we performed dimensionality reduction analysis on the miRNA sequencing data from all 18 individuals. Treat NSCLC patients and treatment-naive NSCLC patient fell into two apparent clusters (Fig. 6b). This indicates substantial differences in miRNA expression associated with health status. Meanwhile, some treated NSCLC patients clustered together with healthy individuals, indicating the similar miRNA expression patterns.

**Fig 6.**
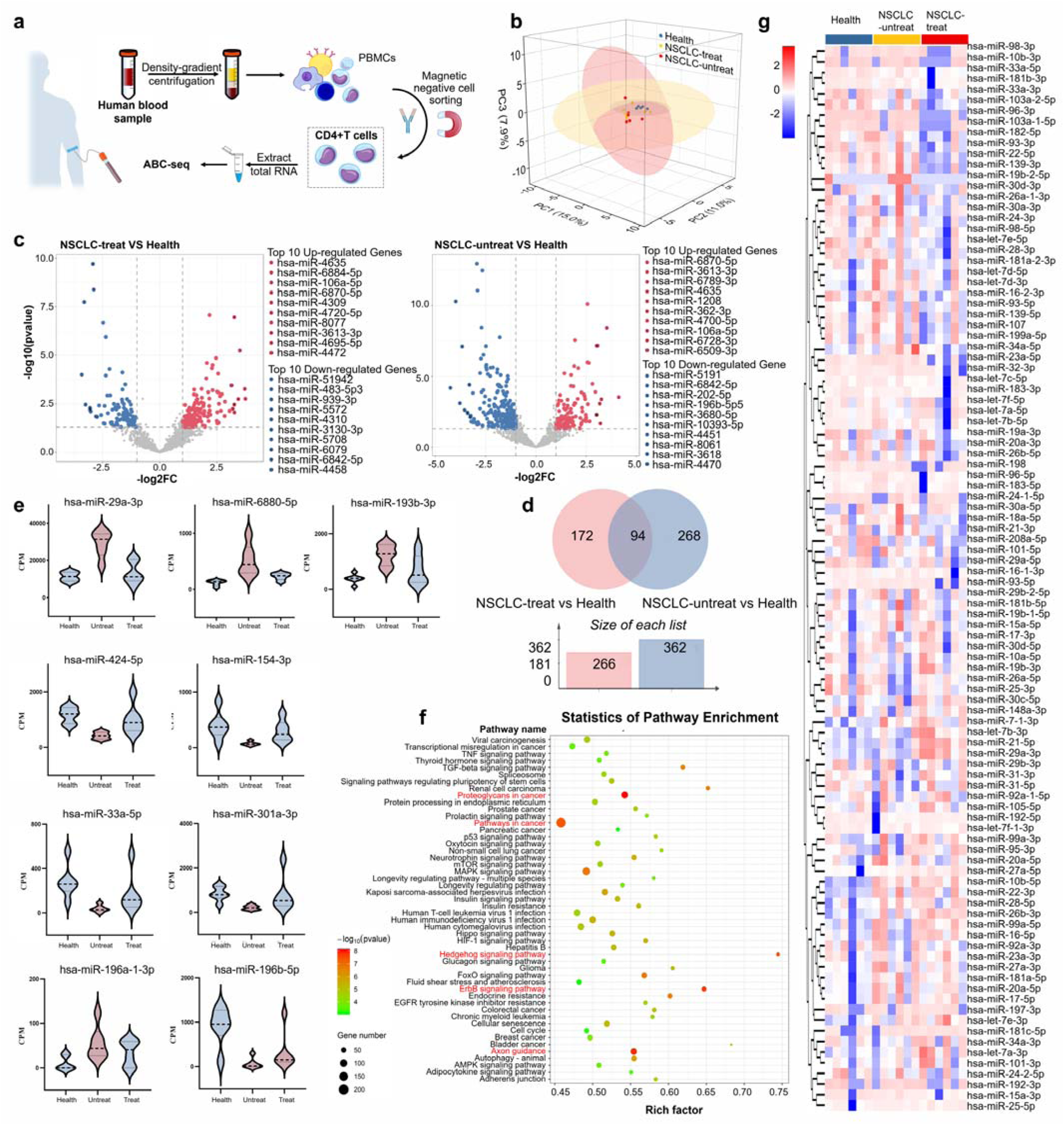
Impact of the slip length on the platelet dynamics. **a** Workflow for ABC-seq analysis of smRNA in CD4+ T cell samples from blood. **b** PCA analysis of samples from healthy individuals (n=6), NSCLC-treated patients (n=6), and NSCLC-untreated patients (n=6). PC1, PC2, and PC3 represent the first, second, and third principal components, respectively. Each point represents a sample, with colors indicating different groups. **c** Volcano plots displaying differential gene expression between two groups. The left plot compares NSCLC-treated patients versus healthy individuals, and the right plot compares NSCLC-untreated patients versus healthy individuals. The x-axis shows the log2 fold change (log2FC), and the y-axis represents the statistical significance of the change in gene expression (-log10(FDR)). Each point represents a specific gene: red points indicate significantly upregulated genes, blue points indicate significantly downregulated genes, and gray points indicate non- significant genes. The rightmost column lists the top 10 genes with the most significant upregulation and downregulation. **d** Venn diagram and bar chart illustrating the overlap and differences in differentially expressed genes between NSCLC-treated versus healthy and NSCLC-untreated versus healthy comparisons (Uniq/Total sRNA). The intersection represents miRNAs highly expressed in patients regardless of treatment status. The pink region on the left indicates miRNAs not differentially expressed when comparing patients to healthy individuals but differentially expressed after treatment. The gray region on the right indicates miRNAs differentially expressed when comparing patients to healthy individuals but not after treatment. **e** Comparative analysis of CPM values for three representative miRNAs selected from the pink, gray, and gray-pink regions of the Venn diagram in panel (d) across the three treatment groups. **f** KEGG pathway enrichment analysis of significantly upregulated miRNA target genes. **g** Heatmap showing clustering of gene expression profiles. The horizontal axis represents sample groups, and the vertical axis represents gene clustering. Samples or genes within the same tree branch are more similar to each other. Different colors indicate relative gene expression levels, with red representing high expression and blue representing low expression.

Differential expression analysis was performed for the treatment group versus the treatment-naive group, the treatment-naive group versus the healthy control group, and the treatment group versus the healthy control group. We identified 41 significantly (Padj < 0.05) upregulated and 26 downregulated miRNAs in the treatment group compared to the untreated group, while there were 44 upregulated and 90 downregulated miRNAs in the untreated versus healthy control comparison. Additionally, we identified 45 significantly (Padj < 0.05) upregulated and 54 downregulated miRNAs in the comparison of the treatment group versus the healthy control group (Fig. 6d). Moreover, comparing the expression of miRNAs from each individual, we found that the miRNA expression patterns in treated patients are more similar to those in healthy individuals than those in untreated patients (Fig. 6c).

Specifically, we identified 172 miRNAs that displayed significant differential expression in untreated cancer patients but normalized to levels observed in healthy individuals following treatment (Fig. 6d, 6g). Among these, miRNAs such as hsa-miR-29a-3p, hsa-miR-6880-5p, and hsa-miR-193b-3p, hsa-miR- 196a-1-3p were notably upregulated in untreated patients and returned to baseline levels post-treatment (Fig. 6e). These findings are consistent with previous research^38–40^, which has reported similar expression patterns for these miRNAs in the context of cancer and immune regulation. The normalization of these miRNAs post-treatment suggests their potential utility as biomarkers for monitoring therapeutic response and disease progression. Conversely, hsa-miR-33a-5p, hsa-miR-154-3p, hsa-miR-424-5p, hsa-miR-301a- 3p exhibited significantly lower expression levels in untreated cancer patients, while their expression levels were restored to levels similar to those in healthy individuals following treatment. On the one hand, this pattern suggests that these miRNAs may play a role in tumor suppression. On the other hand, their downregulation in untreated cancer patients may contribute to tumor progression.

Notably, while previous research has demonstrated that hsa-miR-196b-5p is upregulated in NSCLC tumor tissues, our results indicate that this miRNA is downregulated in CD4+ T cells from peripheral blood in both untreated and treated lung cancer patients^41^. Several potential mechanisms could explain this observation. One of explanation is miRNAs often exhibit tissue-specific expression patterns and regulatory mechanisms. The upregulation of hsa-miR-196b-5p in lung tumor tissues likely reflects its involvement in oncogenic processes within the tumor microenvironment, such as promoting cell proliferation, survival, and metastasis. In contrast, the downregulation in peripheral blood CD4+ T cells suggests that different regulatory factors influence miRNA expression in immune cells.

To understand the biological relevance of the differentially expressed miRNAs, we performed KEGG pathway analysis (Fig. 6f, S9-16). It revealed that miRNAs differentially expressed between untreated and treated cancer patients were significantly enriched in the Neuroactive ligand-receptor interaction, Cell adhesion molecules (CAMs), and Hippo signaling pathway-multiple species pathways. These findings underscore the multifaceted roles of miRNAs in cancer biology, providing insights into their potential as therapeutic targets and biomarkers for treatment efficacy.

## Discussion

High-content analysis of small RNA (smRNA) expression is becoming increasingly critical in life science research. Previous studies have shown that using pooled adapters instead of single adapters can significantly minimize ligation bias^2,24^. However, the lack of effective strategies to minimize pooled adapter dimerization has hindered comprehensive smRNA analysis. In this work, we address this critical limitation by developing ABC-seq. As illustrated in Fig. 4, our findings show that the reduction of adapter dimers significantly enhances the detection of small RNAs. Compared with previous smRNA-seq methods, ABC-seq employs pooled adapters to limit ligation bias while minimizing adapter dimers through both inhibitions of their formation and post-generation excision. This modular design provides researchers with the flexibility to tailor the library preparation process according to specific experimental requirements, allowing for integration with other smRNA-seq methods when necessary.

Future optimizations of ABC-seq may focus on the choice of restriction enzymes. The use of the EcoP15I enzyme, which cleaves at positions 24/25 bp downstream, restricts the RNA region in the 5’ adapter plus the NNN region in the 3’ adapter to a maximum of 25 nucleotides. This limitation confines the current method to sequences within this length. However, future development of enzymes capable of cleaving at longer downstream positions could expand the versatility and application of ABC-seq by accommodating a broader range of sequences.

Overall, ABC-seq offers enhanced accuracy and sensitivity in small RNA detection. ABC-seq is not only a versatile and modular approach designed for low-bias adapter preparation in RNA sequencing but also effectively reduces dimer formation. Beyond its application in smRNA-seq, ABC-seq can serve as a universal upgrade for other short-length RNA profiling, such as ribosome-protected footprints (RPFs) in ribosome profiling^42^, RNA proximity ligation in RNA-RNA interactions^43^, and glycoRNAs on the outer cell surface^44^. In summary, ABC-seq represents a significant advancement in RNA sequencing library preparation methods, offering a novel perspective on high content detection across a wide range of RNA sequencing applications. This method paves the way for more accurate and comprehensive RNA analyses and enriching our understanding of gene expression regulation in physiological and pathological conditions.

## Methods

### Cell culture and clinical samples

The HeLa cell line and the rat heart-derived H9c2 cell line were obtained from Pricella. Both cell lines were cultured in Dulbecco’s Modified Eagle Medium (DMEM, GIBCO), supplemented with 10% fetal bovine serum (Gibco), 1% penicillin-streptomycin (Thermo Fisher). Regular mycoplasma contamination detection was performed. All cells were cultured in 10-cm-diameter dishes and incubated at 37°C in a humidified atmosphere containing 5% CO2. The study of human blood samples was approved by ethics committee of Xi’an Jiaotong University (No. 2021-270) in accordance with the law on human experiments. Informed consent was obtained from all patients.

### Cardiac hypertrophy culture and measurement of cell area

To induce a cardiac hypertrophy model, H9c2 cells were first starved in DMEM without FBS for 24 h, followed by treatment with 0.5 μM Angiotensin II (AngII, Sigma) for 48 h. Post-treatment, the cells were fixed in 4% formaldehyde in PBS at room temperature for 20 min. They were then washed three times with PBS and permeabilized using 0.5% Triton X-100 in PBS for 15 min. Subsequently, the cells were stained with phalloidin (Yeasen, Shanghai, China) and DAPI (Beyotime) to visualize the nuclei. The slides were examined with an Olympus fluorescence microscope at 400x magnification (AX70, Olympus Corporation, Tokyo, Japan). ImageJ software (version 1.8.0) was used to measure the relative cell surface area.

### Collection of FACS and CD4+T cell

Collect 1 mL of peripheral blood from each participant. The study consists of three groups: healthy individuals, non-small cell lung cancer (NSCLC) patients, and NSCLC patients undergoing treatment (chemotherapy and immunotherapy), with six individuals in each group. To prevent clotting, use anticoagulant-treated blood collection tubes (e.g., EDTA).

To facilitate handling, dilute each 1 mL of blood sample with an equal volume of phosphate-buffered saline (PBS). Carefully layer the diluted blood sample over 2 mL of density gradient medium (e.g., Ficoll-Paque) in a 15 mL centrifuge tube. Centrifuge the samples at 400 g for 30 min at room temperature, ensuring that the brake is not applied to maintain layer separation. Following centrifugation, use a pipette to carefully extract the PBMC layer from the interface. Transfer the PBMC layer to a fresh 15 mL tube and wash the PBMCs twice with PBS by centrifuging at 300 g for 10 min, discarding the supernatant, and resuspending the pellet in PBS.

Resuspend the isolated PBMCs in the recommended buffer provided with the MojoSort Human CD4 T Cell Isolation Kit (Biolegend). Add a biotin-conjugated antibody cocktail specific for non-CD4+ cells (e.g., CD8, CD14, CD16, CD19, CD36, CD56, CD123, and CD235a) to the cell suspension. Incubate the suspension at 4°C for 10-15 min. Next, add streptavidin-coated magnetic beads to the cell suspension and incubate at 4°C for an additional 10-15 min. Place the tube in a magnetic separator for the recommended duration (typically 5-10 min) to allow the non-CD4+ cells to bind to the beads and remain in the tube.

Carefully collect the supernatant containing the enriched CD4+ T cells into a new tube. Count the cells to ensure there is an adequate number for RNA extraction.

### Total RNA and small RNA Extraction

For HeLa and H9c2 cells, culture them until they reach a maximum of 90% confluence. Wash the cells twice with PBS and remove the supernatant by pipetting. 1 mL of TRIzol (Thermo Fisher) was added to the dish to lysis cells. If extracting small RNA, instead of TRIzol, add 1 mL of RNAiso for Small RNA (RSR, Takara). For the enriched CD4+ T cells, resuspend the cell pellet in 1 mL of TRIzol reagent or Small RNAiso per 0.5-5 million cells, ensuring complete lysis by pipetting up and down or briefly vortexing. Incubate the lysed cells in TRIzol for 5 min at room temperature to allow complete dissociation of nucleoprotein complexes.

Add 0.2 mL of chloroform per 1 mL of TRIzol or RSR reagent, securely cap the tube, shake it vigorously by hand for 15 seconds, and incubate the mixture at room temperature for 2-3 min. Centrifuge the tube at 12,000 g for 15 min at 4°C to separate the mixture into three phases: a clear upper aqueous phase containing RNA, an interphase, and a red lower organic phase. Carefully transfer the clear aqueous phase to a new tube, avoiding the interphase and organic phase.

Add 0.5 mL of isopropanol per 1 mL of TRIzol or RSR reagent used initially, mix the sample by inverting the tube several times, and incubate the mixture at room temperature for 10 min to precipitate the RNA. Centrifuge at 12,000 g for 10 min at 4°C to pellet the RNA. Remove the supernatant carefully without disturbing the RNA pellet. Wash the RNA pellet by adding 1 mL of 75% ethanol, vortex briefly, and centrifuge at 7,500 g for 5 min at 4°C. Remove the ethanol carefully and air-dry the RNA pellet for 5- 10 min, ensuring the pellet does not dry completely as it will be difficult to dissolve.

Dissolve the RNA pellet in an appropriate volume of RNase-free water. Measure the RNA concentration using a spectrophotometer or NanoDrop instrument, and assess RNA quality by running an aliquot on an agarose gel or using a Bioanalyzer to check the integrity of the RNA.

### Library preparation protocol of ABC-seq

ssRNA 3’ random ligation : The RNA sample was mixed with 5 pmol of 3’ adapter pool, 1 μL of 10X T4 ligase reaction buffer, 2 μL of 50% PEG8000, 40 U of T4 RNA Ligase 2, truncated K227Q (200 U/μL, NEB), 20 U of Recombinant RNase Inhibitor (Takara), and RNase-free water to a final volume of 10 μl. The mixture was incubated at 30°C for 6 h.

Block adapter by DNA polymerization: To the ligation product, add 1 μL of 20 μM RT primer. Incubate the mixture sequentially at 75°C for 5 min, 45°C for 15 min, 25°C for 15 min, and 4°C for 5 min. Next, add 5 U of phi29 DNA Polymerase (10 U/μL, NEB), 0.5 μL of 10 mM dNTPs, and RNase-free water to achieve a final reaction volume of 15 μL. The sample was incubated at 30°C for 1 h.

ssRNA 5’ random ligation: Mix the ligation product with 20 pmol of 5’ adapter pool, 1 μL of 10X T4 ligase reaction buffer, 2 μL of 10 mM ATP, and 30 U of T4 RNA Ligase 1 (30 U/mL, NEB). The sample was incubated at 37°C for 1 h.

Reverse transcription: All the ligation products were used as templates for RT in a final 30 μl reaction volume containing 8 U of Bst 3.0 DNA Polymerase (8 U/μL, NEB), 1X RT reaction buffer, 1 μL of 10 mM dNTP, and 40 U of RRI. The reaction was performed at 55°C for 1 h and 85°C for 20 min.

RE treatment: Mix the 3’ adapter and RT primer at an equimolar ratio in 1X T4 RNA Ligase reaction buffer in an appropriate volume. Incubate this mixture at 37°C for 15 min. Then, add 0.2 pmol of the preincubated mixture to the RT reaction and follow with the addition of 25 U of EcoP15I (10 U/μL, NEB). Incubate the RT reaction at 37°C for 2 h and then at 85°C for 20 min.

PCR amplification: Amplify the RT product in a 20 μl reaction volume containing 1 μl of the RT reaction product, 1 U of Hot Start Taq DNA Polymerase (5 U/μL, NEB), 1X PCR buffer, 5 pmol of the RPI primer, 5 pmol of the index RPI primer, 1 μL of 10X T4 ligase reaction buffer, and RNase-free water.

Heat the PCR mixture to 94°C for 2 min, followed by 15–18 cycles (for 200 ng input small RNA) of 94°C for 10 s, 62°C for 30 s, and 68°C for 15 s, with a final extension at 68°C for 5 min.

Sequencing: Small RNA libraries were sequenced using either the Illumina NovaSeq-6000 or the DNBSEQ-T7 sequencer.

The oligo sequences for the 3’ adapter, RT primer, 5’ adapter, and PCR primer used in the construction of the small RNA library are listed in Supplementary Table 1.

### Data analysis

Raw data (raw reads) in FASTQ format were first processed using in-house Perl scripts. During this step, clean data (clean reads) were obtained by removing reads containing adapters, reads with poly-N sequences, and low-quality reads. Additionally, reads shorter than 16 nt or longer than 40 nt were trimmed and removed. Quality metrics such as Q20, Q30, GC content, and sequence duplication levels of the clean data were calculated. All downstream analyses were conducted using high-quality clean data.

Using Bowtie, the clean reads were aligned to the Silva, GtRNAdb, piRBase, Rfam, and Repbase databases to filter out ribosomal RNA (rRNA), transfer RNA (tRNA), PIWI-interacting RNA (piRNA), small nuclear RNA (snRNA), small nucleolar RNA (snoRNA), other non-coding RNAs (ncRNAs), and repeats. The remaining reads were then used to detect known miRNAs and predict novel miRNAs by comparing them with the genome and known miRNAs from miRBase. Novel miRNA secondary structure prediction was performed using Randfold and RNAfold tools.

Gene function was annotated based on the following databases: Nr (NCBI non-redundant protein sequences), Pfam (Protein family), KOG/COG (Clusters of Orthologous Groups of proteins), Swiss-Prot (A manually annotated and reviewed protein sequence database), KEGG (KEGG Ortholog database), and GO (Gene Ontology). Gene Ontology (GO) enrichment analysis of the differentially expressed genes (DEGs) was implemented by the ClusterProfiler R packages based Wallenius non-central hyper- geometric distribution. KEGG (Kanehisa et al., 2008) was used to test the statistical enrichment of differential expression genes in KEGG pathways.

miRNA expression levels were estimated for each sample by mapping sRNAs back onto the precursor sequences. Read counts for each miRNA were obtained from the mapping results. For samples with biological replicates, differential expression analysis between two conditions/groups was performed using the DESeq2 R package (1.10.1). DESeq2 provides statistical routines for determining differential expression in digital miRNA expression data using a model based on the negative binomial distribution. The resulting P-values were adjusted using the Benjamini and Hochberg approach to control the false discovery rate. miRNAs with |log2(FC)| ≥ 1.00 and FDR ≤ 0.05 were considered differentially expressed. Differential expression analysis was performed using BMKCloud (www.biocloud.net).

### Statistical information

All data are presented as mean ± standard deviation (s.d.). Statistical analyses were performed using GraphPad Prism software. The significance between two groups was assessed using Student’s t-test with unpaired two-sided comparisons. For comparisons among multiple groups, one-way analysis of variance (ANOVA) with Tukey’s multiple comparisons test or two-way ANOVA with Bonferroni’s multiple comparisons test was employed. Prior to differential gene expression analysis, differential expression analysis of samples was performed using edgeR. P-values were adjusted using the q-value method (Storey et al., 2003). A threshold of |log2(FC)| ≥ 1.00 and FDR ≤ 0.01 was set for significant differential expression.

## Data Availability

The sequencing data generated in this study have been deposited at Gene Expression Omnibus (GE0) with accession number GSE274194.

## Acknowledgements

This work was supported by the National Natural Science Foundation of China (No. 22125404), the Natural Science Basic Research Program of Shaanxi (2023-JC-JQ-13), the Innovation Capability Support Program of Shaanxi (2023-CX-TD-62), the Fundamental Research Funds for the Central Universities and the “Young Talent Support Plan” of Xi’an Jiaotong University.

## Ethics declarations

### Contributions

H.H.Y., F.C., and Y.X.Z. designed the experiments and wrote the manuscript. H.H.Y. and M.Y.Y. performed the majority of experiments and analysed the data. H.H.Y., Y.Y.G., and Y.T.S. created the schematics and data visualizations. K.C. provided the HeLa and H9C2 cell lines and guided the cell culture procedures. D.X.Z. supplied the cell and clinical samples and conducted the clinical data analysis.

