## Supplemental files for "ABC-seq expands small RNAs content with randomized adapter pools operation"

**Supplement Table1:**

| Oligo Name | Sequences |
|---|---|
| miR-545 | UCAGCAAACAUUUAUUGUGUGC |
| miR-145 | GUCCAGUUUUCCCAGGAAUCCCU |
| miR-191 | CAACGGAAUCCCAAAAGCAGCUG |
| miR-31 | AGGCAAGAUGCUGGCAUAGCU |
| miR-125b | UCCCUGAGACCCUAACUUGUGA |
| 3' randomized adapter | 5'-p-NNNCTGCTGTGGAATTCTCGGGTGCCAAGG-Cy5-3' |
| 5' randomized adapter | GTTCAGAGTTCTACA*rGrUrCrCrGrArCrGrArUrCrArGrNrNrNrN* |
| RT primer | CCTTGGCACCCGAGAATTCCACAGCAG |
| NNN RT primer | CCTTGGCACCCGAGAATTCCACAGCAGNNN |
| RP1 primer | AATGATACGGCGACCACCGAGATCTACACGTTCAGAGTTCTACAGTCCGA |
| INDEX-RPI1 | CAAGCAGAAGACGGCATACGAGATCGTGATGTGACTGGAGTTCCTTGGCACCCGAGAATTCCA |
| INDEX-RPI2 | CAAGCAGAAGACGGCATACGAGATACATCGGTGACTGGAGTTCCTTGGCACCCGAGAATTCCA |
| INDEX-RPI3 | CAAGCAGAAGACGGCATACGAGATGCCTAAGTGACTGGAGTTCCTTGGCACCCGAGAATTCCA |
| INDEX-RPI4 | CAAGCAGAAGACGGCATACGAGATTGGTATGTGACTGGAGTTCCTTGGCACCCGAGAATTCCA |
| INDEX-RPI5 | CAAGCAGAAGACGGCATACGAGATCACTGTGTGACTGGAGTTCCTTGGCACCCGAGAATTCCA |

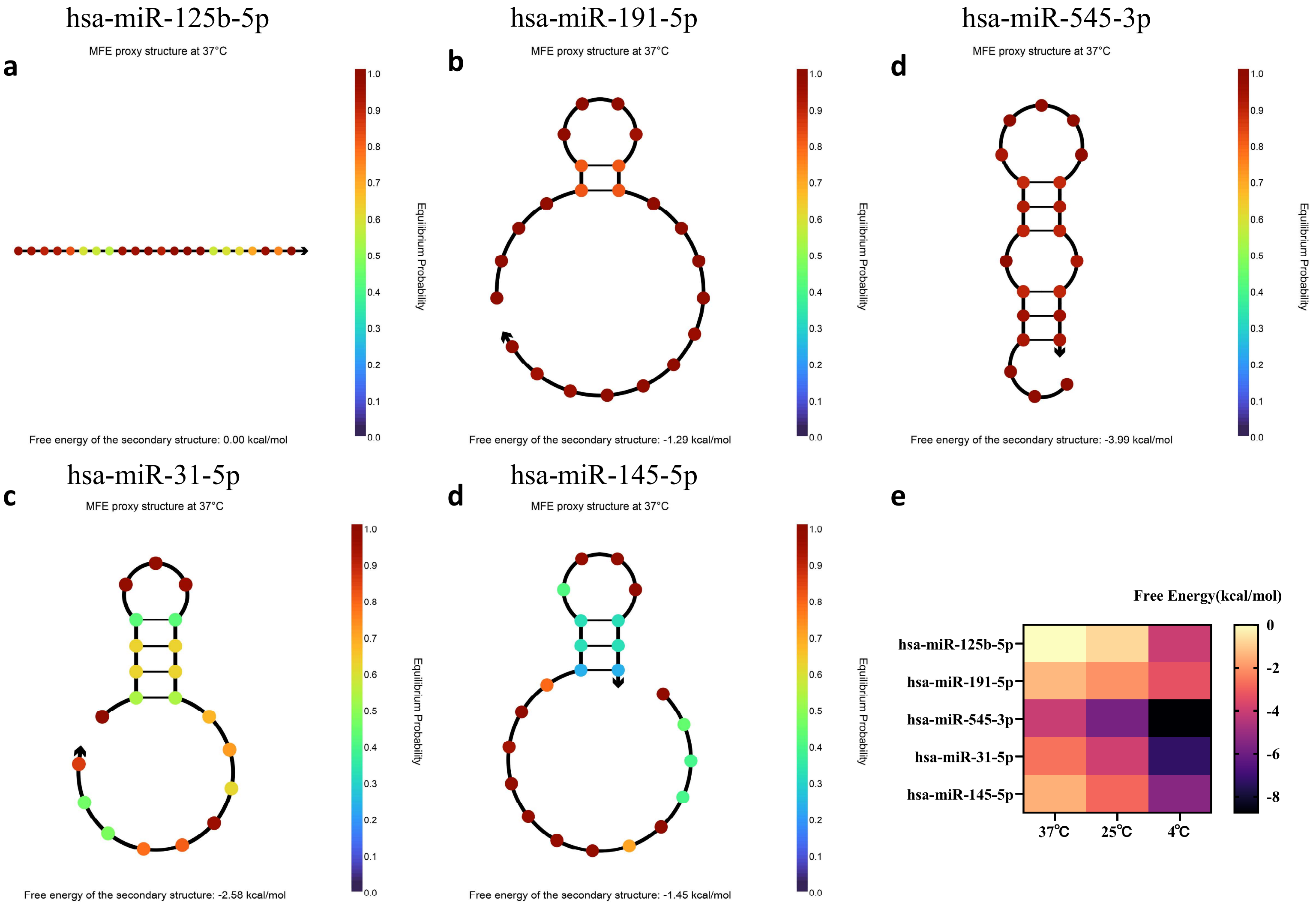


**Figure S1. Secondary Structures and Free Energy of Five miRNAs at Various Temperatures. (a)-(b)** Secondary Structure of hsa-miR-125b-5p, hsa-miR-191-5p, hsa-miR-545-3p, hsa-miR-31-5p and hsa-miR-145-5p at 37°C. **(e)** Free Energy of Various miRNAs at 4°C, 25°C, and 37°C

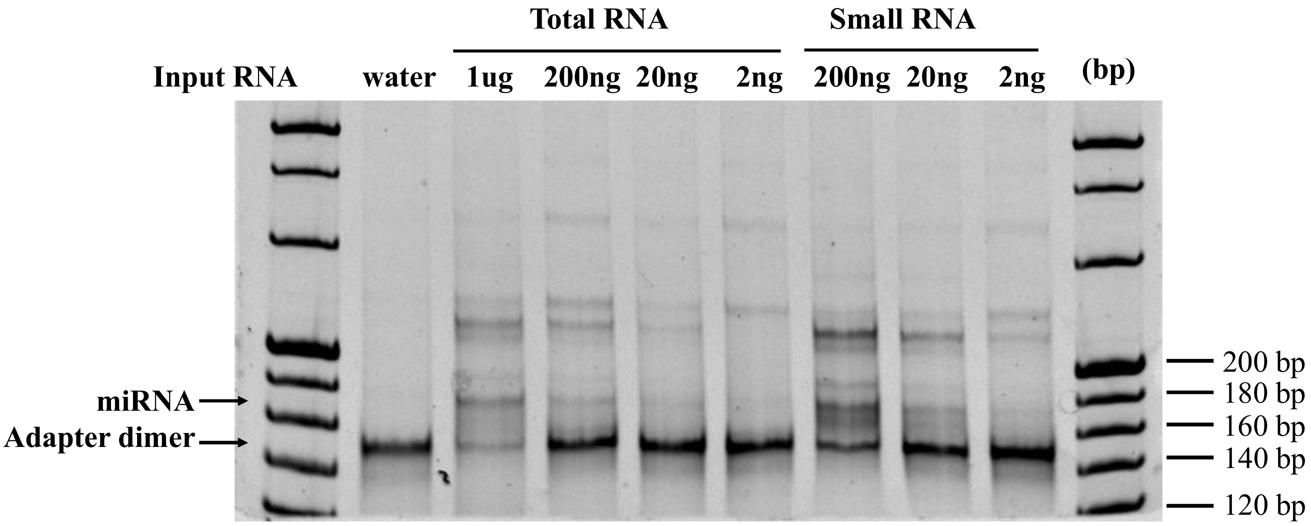


**Figure S2. Sensitivity Assessment of ABC-seq with Varying Input Amounts of Total and Small RNA.** Gel electrophoresis demonstrates ABC-seq sensitivity by comparing the product yields with input RNA amounts ranging from 1 μg to 2 ng for both total and small RNA samples. The 150 bp and 170 bp bands correspond to adapter dimers and miRNAs (20 nt), respectively.

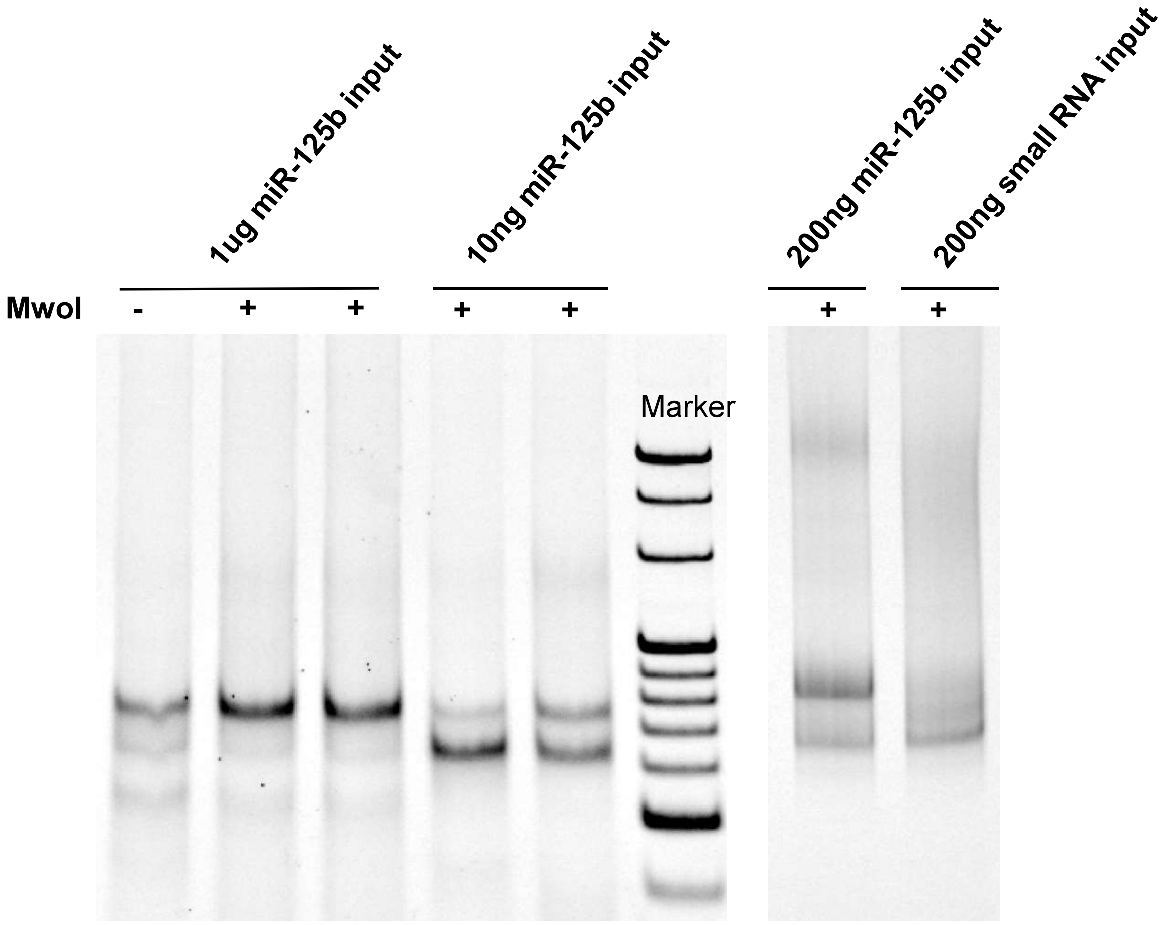


**Figure S3. The performance of Library Construction Efficiency Using Type IIS RE (MwoI) for AC-seq with 1 µg miR-125b, 10 ng miR-125b, and 1 µg Total RNA.** The results show a significant increase in library construction efficiency of miR-125b after enzymatic cleavage.

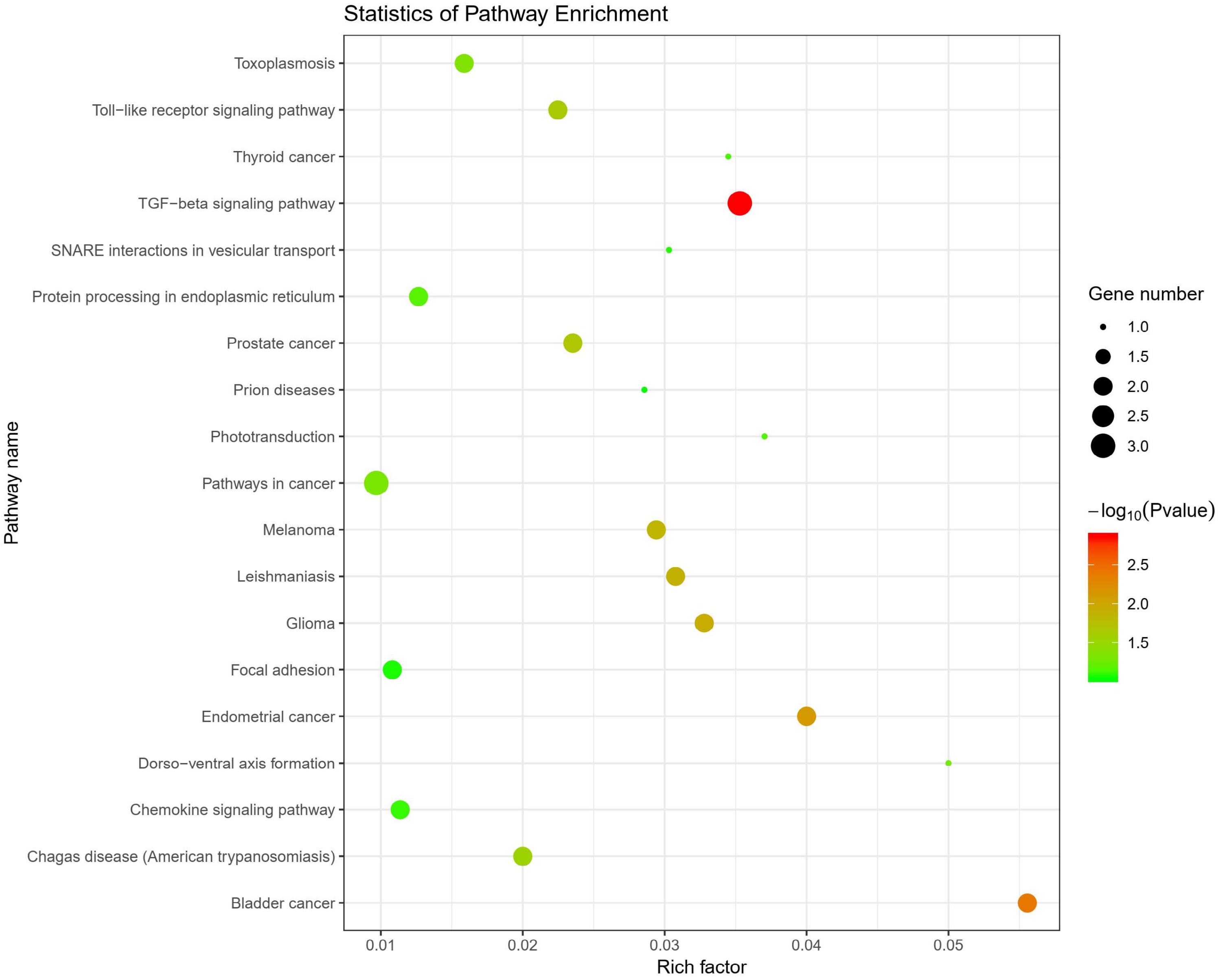


**Figure S4. GO Pathway Analysis Reveals Differential miRNA Expression between Cardiomyocytes and Hypertrophic Cardiomyocytes Using ABC-seq.** The bubble plot illustrates significant upregulation of miRNAs (fold change > 2, $p < 0.05$) and their associated target pathways. Data represent n=2 biological replicates.

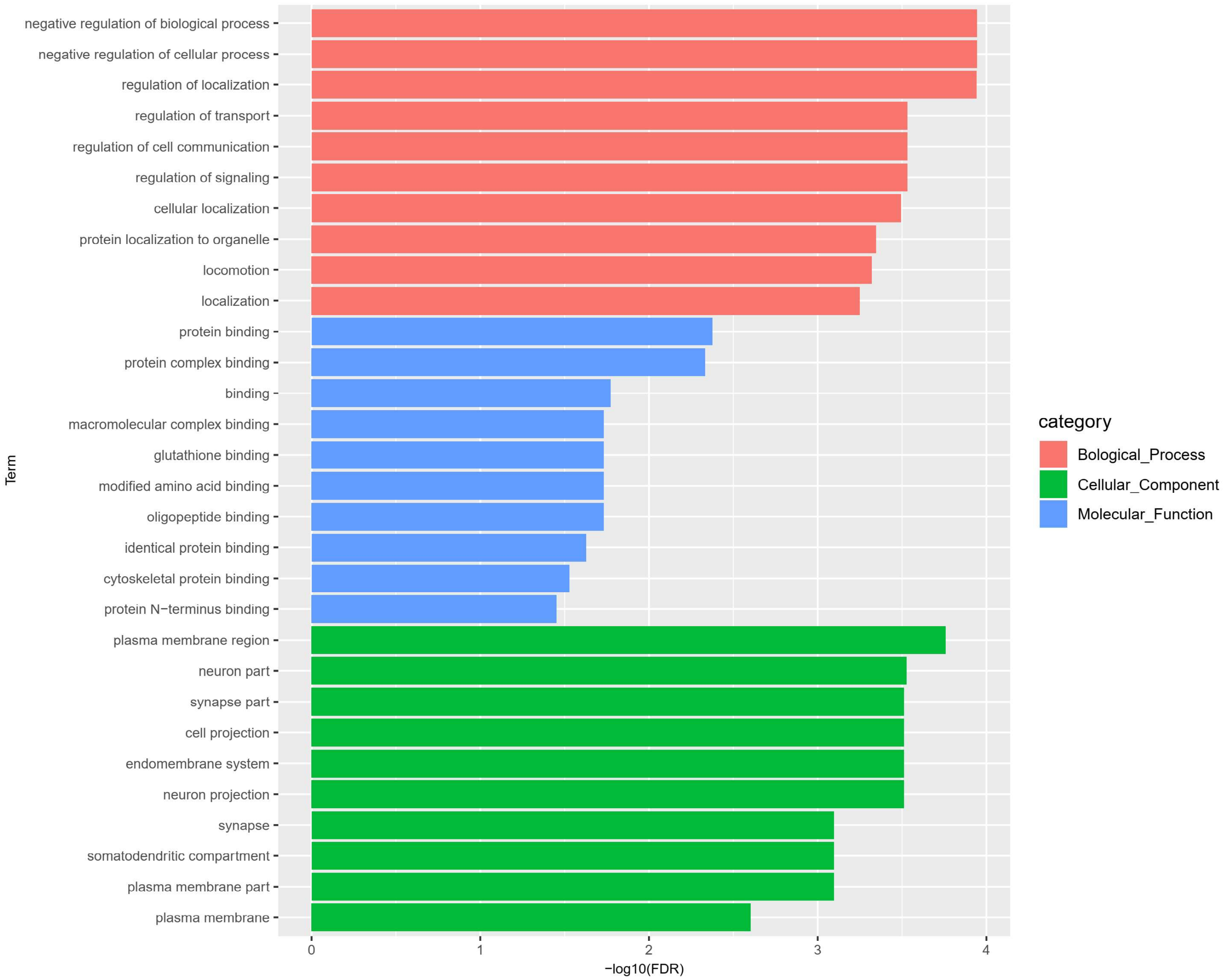


**Figure S5. GO Category Analysis of Differential miRNA Expression between Cardiomyocytes and Hypertrophic Cardiomyocytes Using ABC-seq.** Bar chart illustrating significant upregulated GO categories targeted by miRNAs in cardiomyocytes and hypertrophic cardiomyocytes. Data represent n=2 biological replicates.

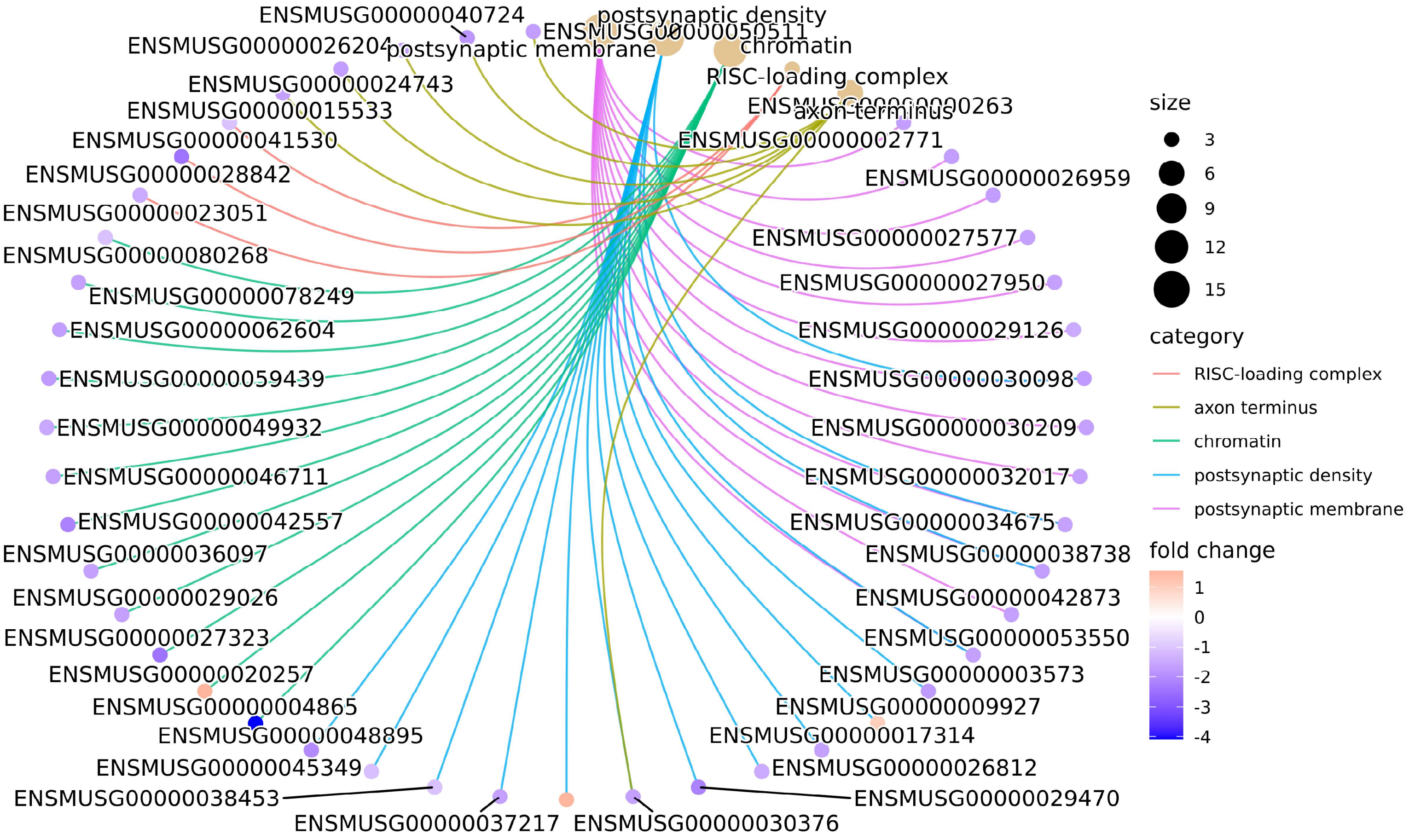


**Figure S6. KEGG Pathway Enrichment in Cardiomyocytes and Hypertrophic Cardiomyocytes.** KEGG pathway enrichment analysis showing enriched pathways in cardiomyocytes and hypertrophic cardiomyocytes identified using ABC-seq. Data represent n=2 biological replicates..

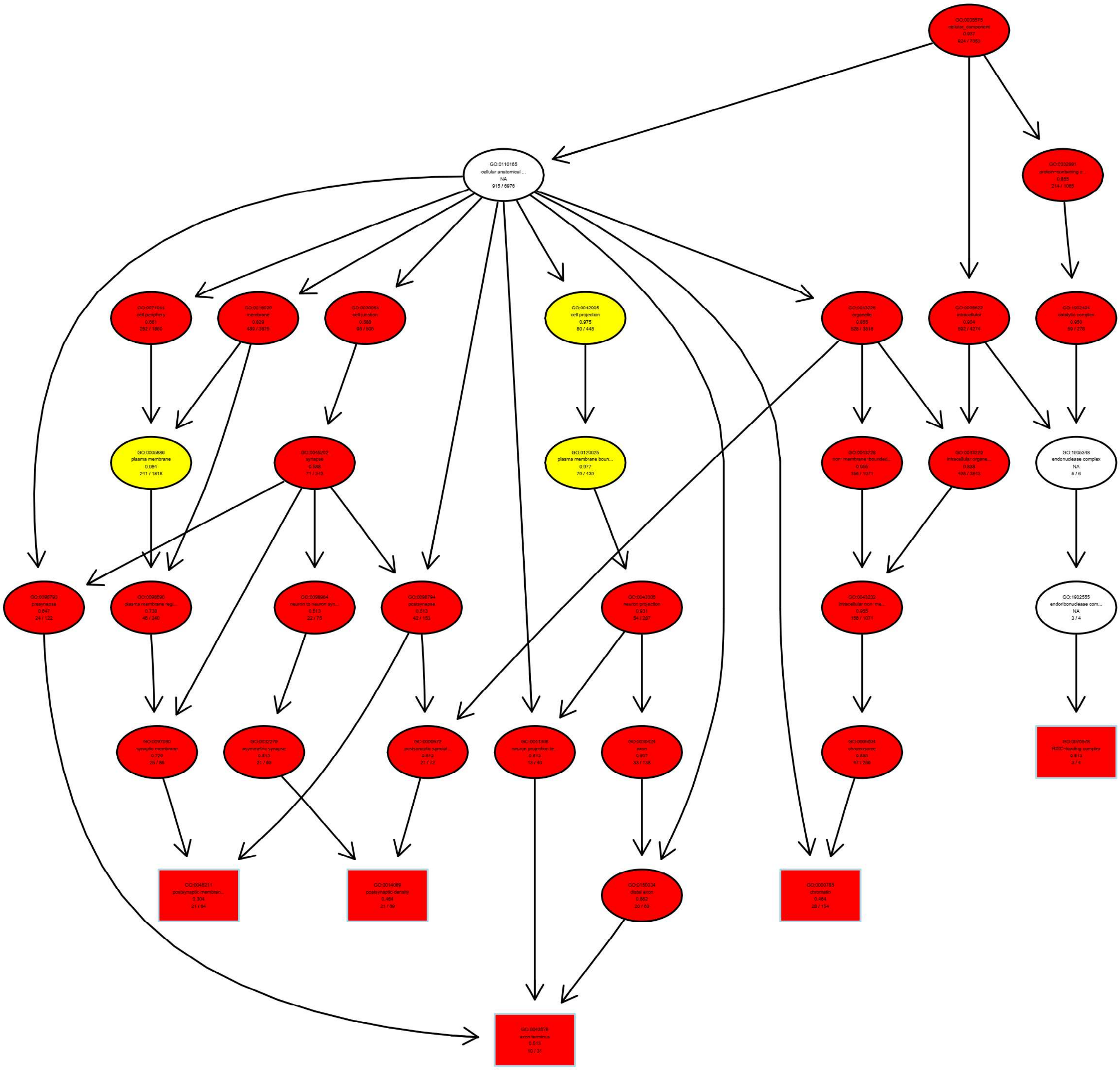


**Figure S7. TopGO Analysis of Cellular Component GO Terms in Cardiomyocytes and Hypertrophic Cardiomyocytes.** TopGO analysis of Cellular Component GO terms depicting enriched categories in cardiomyocytes and hypertrophic cardiomyocytes. Data represent n=2 biological replicates.

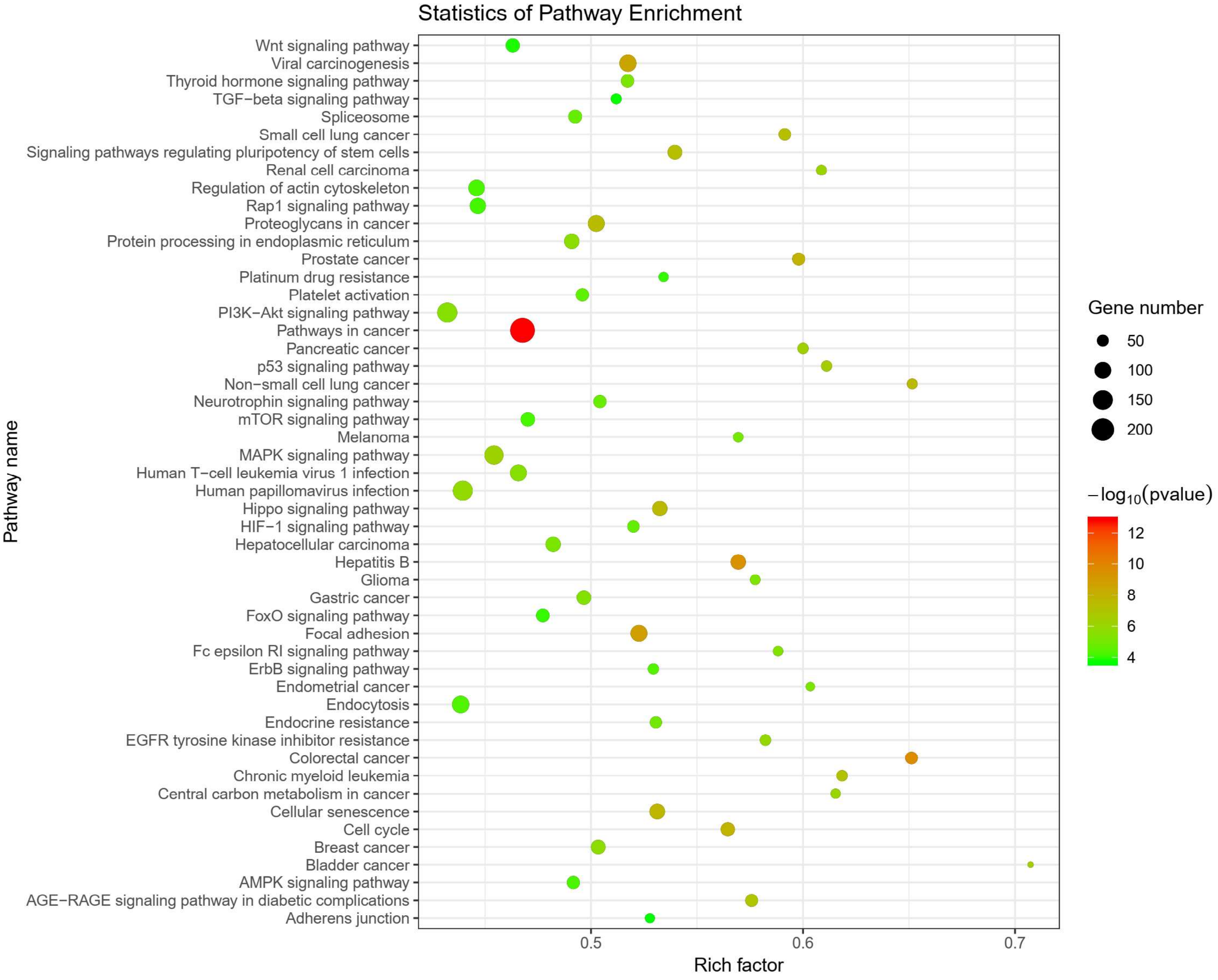


**Figure S8. GO Pathway Analysis Reveals Differential miRNA Expression between Treat and Untreat NSCLC patients Using ABC-seq.** Bubble plot illustrating significant downregulation of miRNAs (fold change > 2, $p < 0.05$) and their associated target pathways in treated versus untreated NSCLC patients using ABC-seq. Data represent n=6 biological replicates.

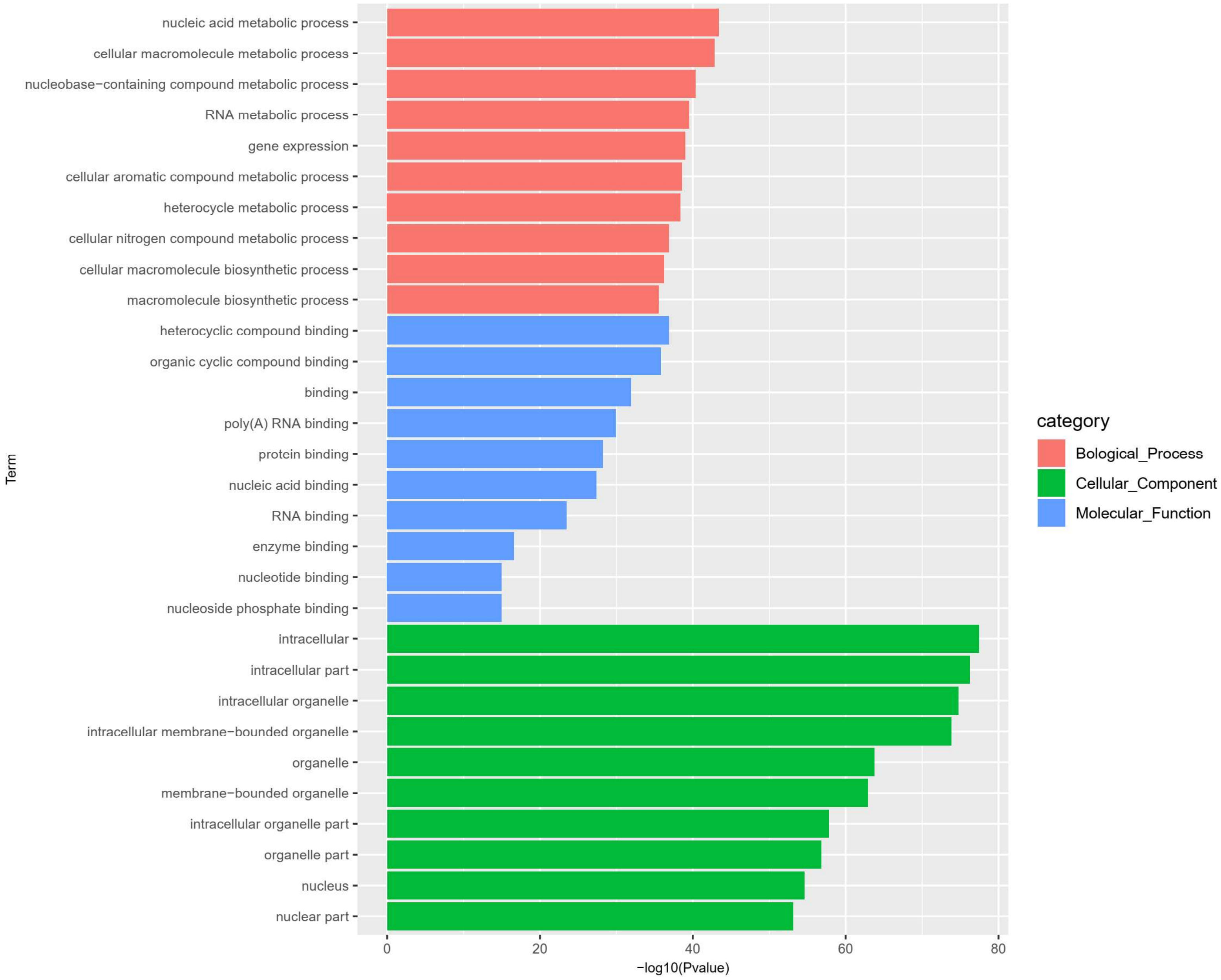


**Figure S9. GO Category Analysis of Differential miRNA Expression between Treat and Untreat NSCLC patients Using ABC-seq.** Bar chart illustrating significant downregulated GO categories targeted by miRNAs in Treat and Untreat NSCLC patients. Data represent n=6 biological replicates.

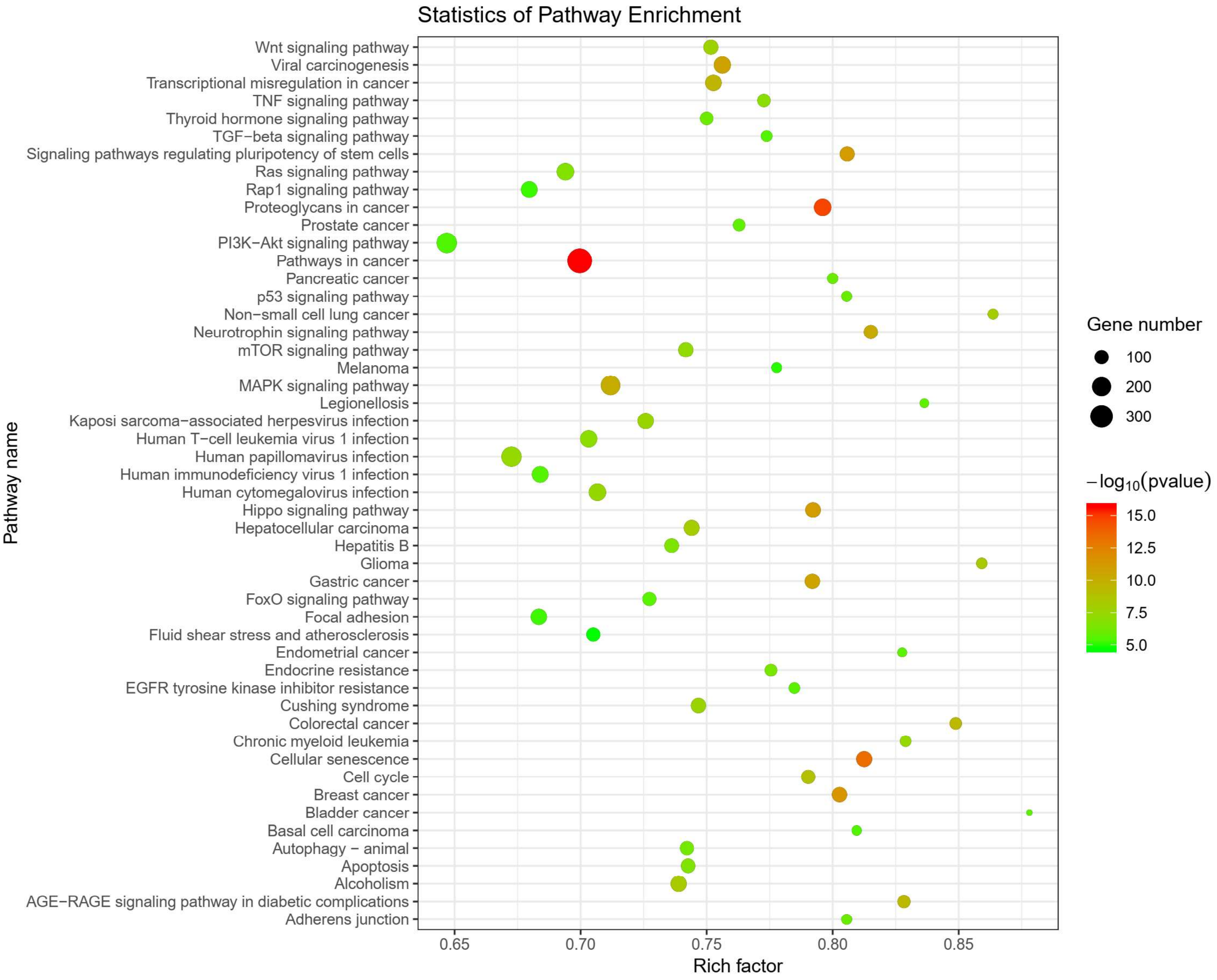


**Figure S10. GO Pathway Analysis Reveals Differential miRNA Expression between Treat and Untreat NSCLC patients Using ABC-seq.** Bubble plot illustrating significant upregulation of miRNAs (fold change > 2, $p < 0.05$) and their associated target pathways in treated versus untreated NSCLC patients using ABC-seq. Data represent n=6 biological replicates.

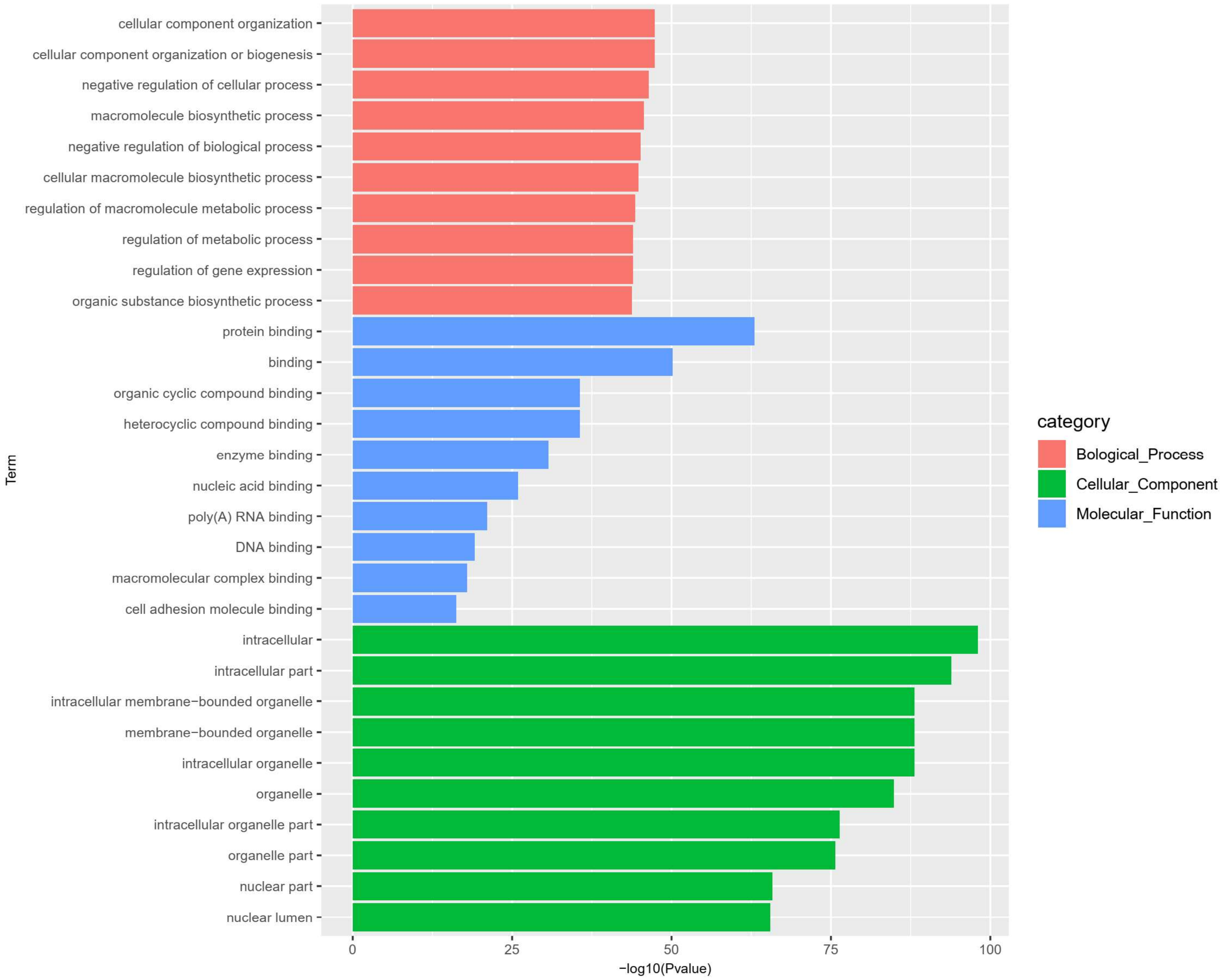


**Figure S11. GO Category Analysis of Differential miRNA Expression between Treat and Untreat NSCLC patients Using ABC-seq.** Bar chart illustrating significant upregulated GO categories targeted by miRNAs in Treat and Untreat NSCLC patients. Data represent n=6 biological replicates.

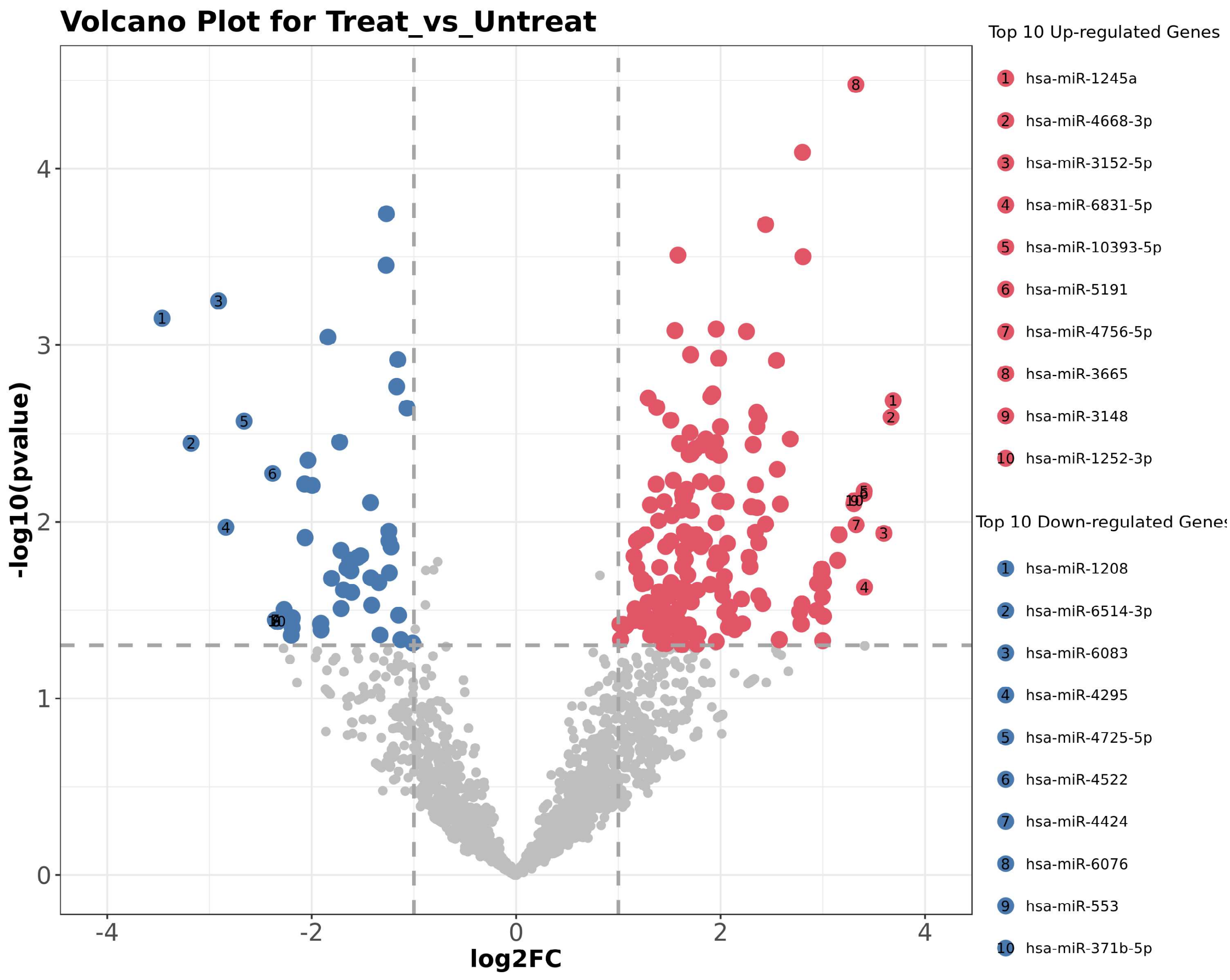


**Figure S12. Volcano Plot of Differentially Expressed Genes in Treated and Untreated NSCLC Patients.** Volcano plot depicting differential expression of genes between treated and untreated NSCLC patients. Genes with a fold change > 2 and p < 0.05 are highlighted as significantly differentially expressed.

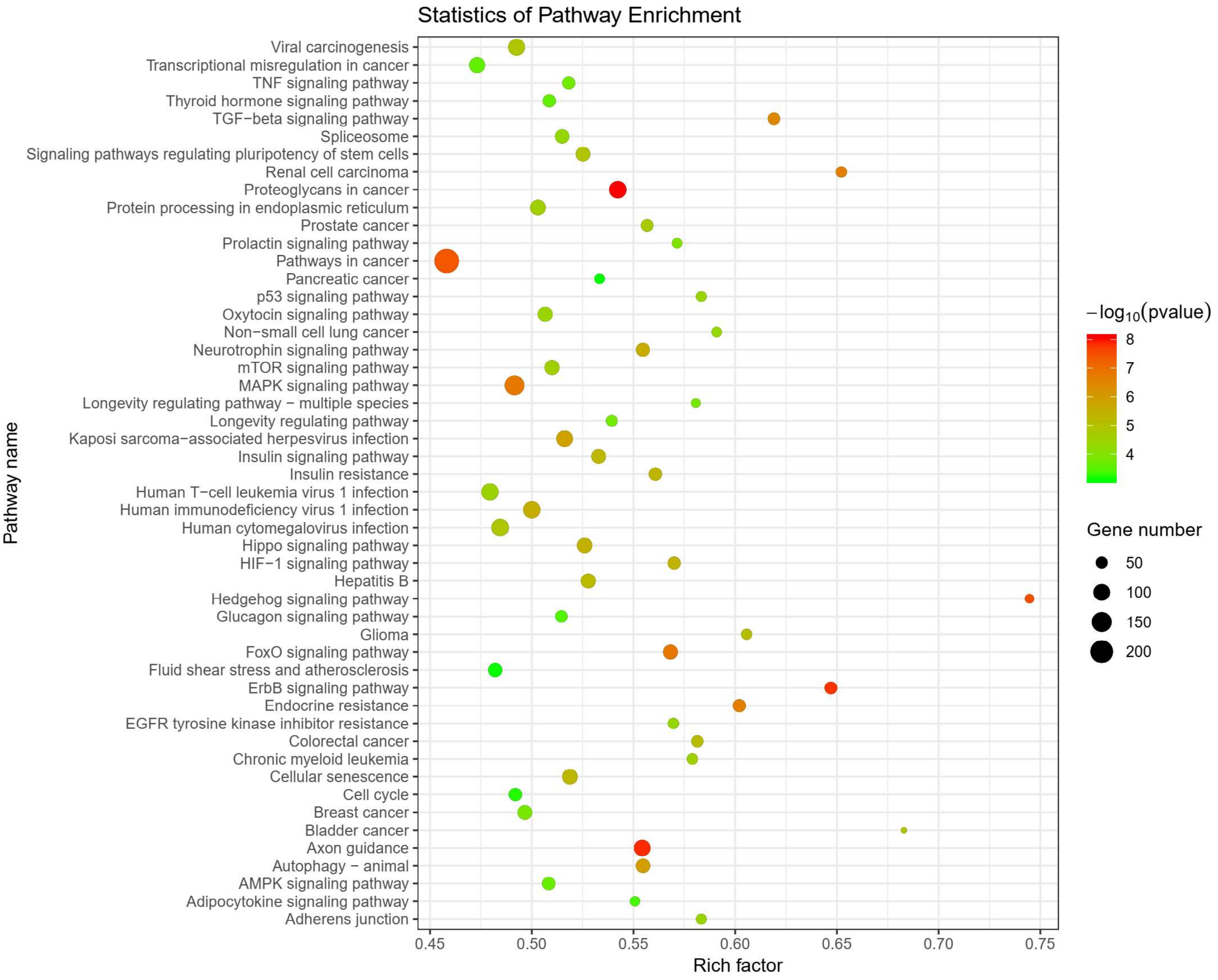


**Figure S13. GO Pathway Analysis Reveals Differential miRNA Expression between NSCLC patients and Health individuals Using ABC-seq.** Bubble plot illustrating significant downregulation of miRNAs (fold change > 2, $p < 0.05$) and their associated target pathways in NSCLC patients versus Health individuals using ABC-seq. Data represent n=6 biological replicates.

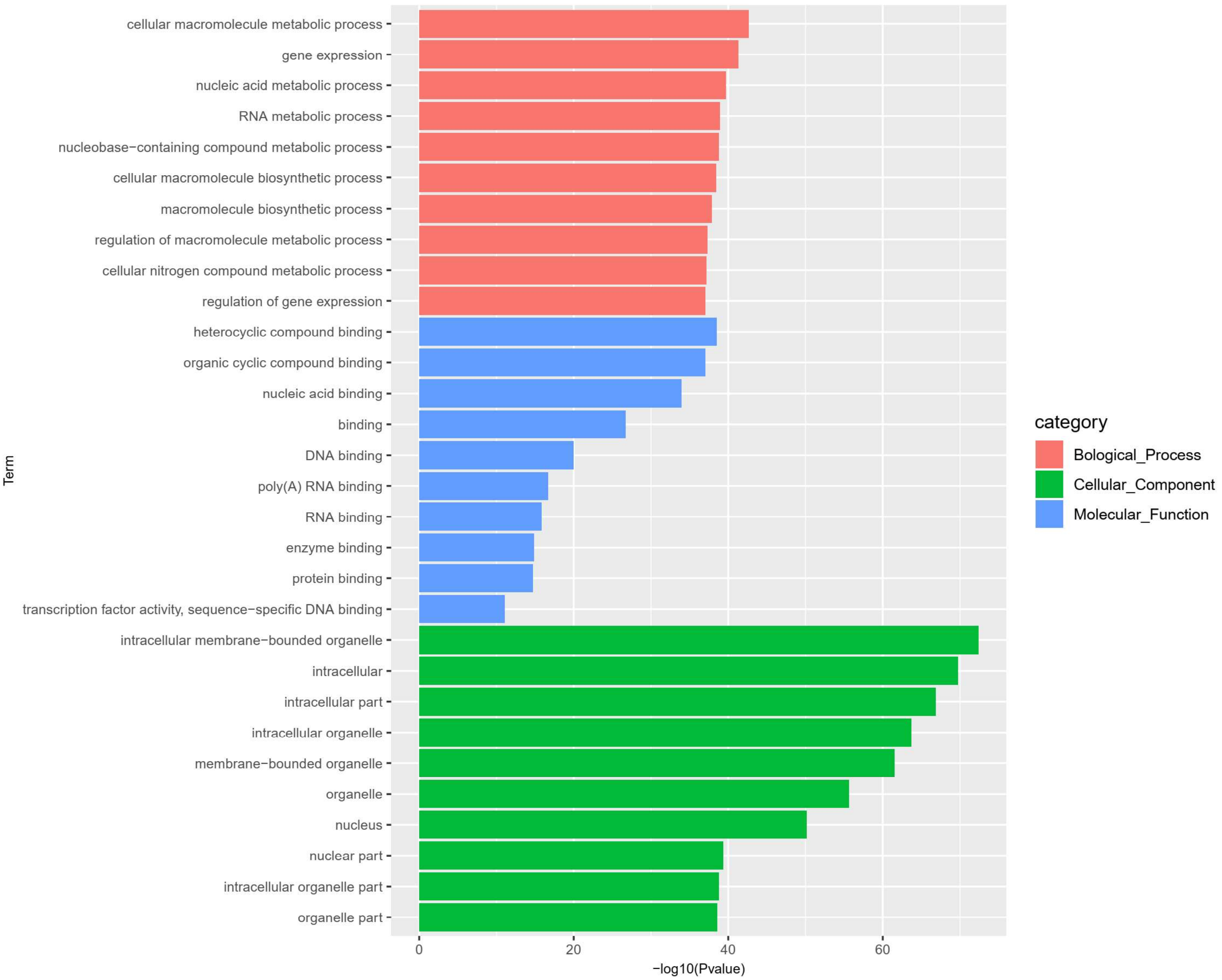


**Figure S14. GO Category Analysis of Differential miRNA Expression between NSCLC Patients and Healthy Individuals Using ABC-seq.** Bar chart illustrating significant downregulated GO categories targeted by miRNAs in NSCLC patients compared to healthy individuals using ABC-seq. Data represent n=6 biological replicates.

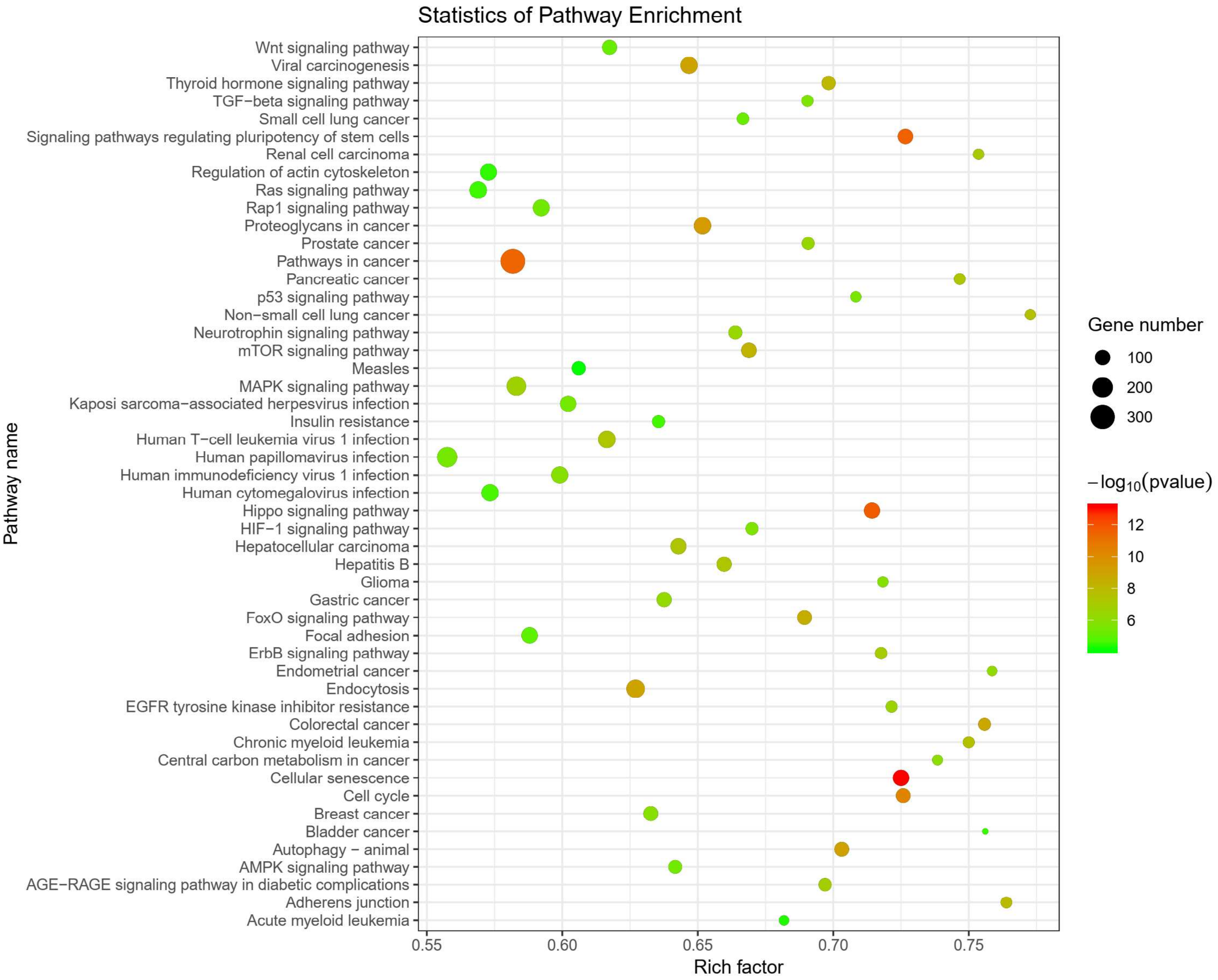


**Figure S15. GO Pathway Analysis Reveals Differential miRNA Expression between NSCLC patients and Health individuals Using ABC-seq.** Bubble plot illustrating significant upregulation of miRNAs (fold change > 2, p < 0.05) and their associated target pathways in NSCLC patients versus Health individuals using ABC-seq. Data represent n=6 biological replicates.

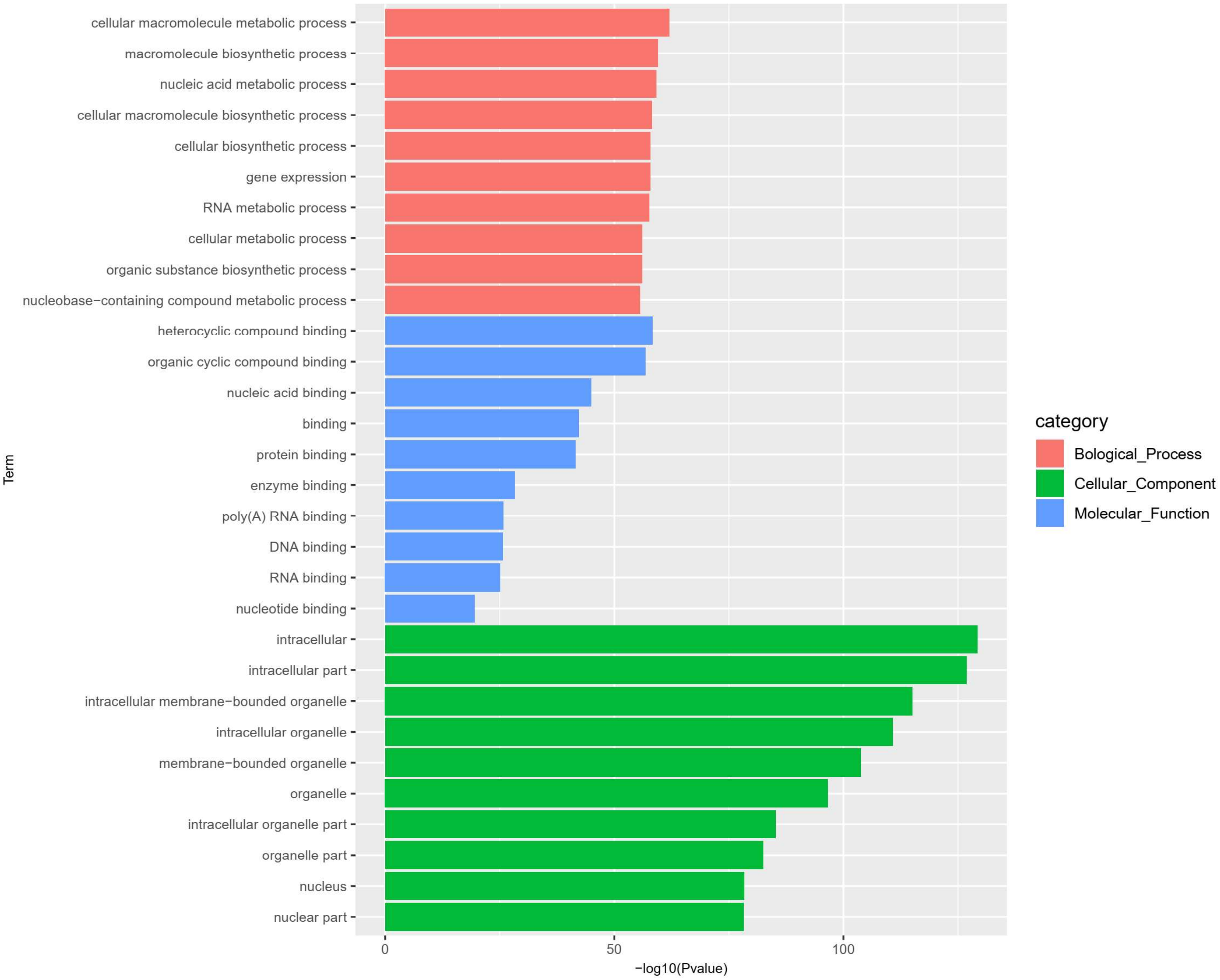


**Figure S16. GO Category Analysis of Differential miRNA Expression between NSCLC Patients and Healthy Individuals Using ABC-seq.** Bar chart illustrating significant upregulated GO categories targeted by miRNAs in NSCLC patients compared to healthy individuals using ABC-seq. Data represent n=6 biological replicates.
